# Deoxyuridine-rich cytoplasmic DNA antagonizes STING-dependent innate immune responses and sensitizes resistant tumors to anti-PD-L1 therapy

**DOI:** 10.1101/2024.04.04.588079

**Authors:** Pinakin Pandya, Frank P. Vendetti, Joseph El-Ghoubaira, Sudipta Pathak, Joshua J. Deppas, Reyna Jones, Anthony V. Columbus, Yunqi Zhang, Daniel Ivanov, Ziyu Huang, Kate M. MacDonald, Shane M. Harding, Raquel Buj, Katherine M. Aird, Jan H. Beumer, Robert W. Sobol, Christopher J. Bakkenist

## Abstract

DNA damage and cytoplasmic DNA induce type-1 interferon (IFN-1) and potentiate responses to immune checkpoint inhibitors. Our prior work found that inhibitors of the DNA damage response kinase ATR (ATRi) induce IFN-1 and deoxyuridine (dU) incorporation by DNA polymerases, akin to antimetabolites. Whether and how dU incorporation is required for ATRi-induced IFN-1 signaling is not known. Here, we show that ATRi-dependent IFN-1 responses require uracil DNA glycosylase (UNG)-initiated base excision repair and STING. Quantitative analyses of nine distinct nucleosides reveals that ATRi induce dU incorporation more rapidly in UNG wild-type than knockout cells, and that induction of IFN-1 is associated with futile cycles of repair. While ATRi induce similar numbers of micronuclei in UNG wild-type and knockout cells, dU containing micronuclei and cytoplasmic DNA are increased in knockout cells. Surprisingly, DNA fragments containing dU block STING-dependent induction of IFN-1, MHC-1, and PD-L1. Furthermore, UNG knockout sensitizes cells to IFN-γ *in vitro*, and potentiates responses to anti-PD-L1 in resistant tumors *in vivo*. These data demonstrate an unexpected and specific role for dU-rich DNA in suppressing STING-dependent IFN-1 responses, and show that UNG-deficient tumors have a heightened response to immune checkpoint inhibitors.

**STATEMENT OF SIGNIFICANCE:** Antimetabolites disrupt nucleotide pools and increase dU incorporation by DNA polymerases. We show that unrepaired dU potentiates responses to checkpoint inhibitors in mouse models of cancer. Patients with low tumor UNG may respond to antimetabolites combined with checkpoint inhibitors, and patients with high tumor UNG may respond to UNG inhibitors combined with checkpoint inhibitors.

## INTRODUCTION

Uracil (U) is a nucleobase lesion in DNA that results from cytosine deamination and from the incorporation of deoxyuridine (dU) into DNA by replicative and repair DNA polymerases (1–3). Widely used standard-of-care antimetabolites inhibit thymidylate synthase, increase the ratio of free dUTP:TTP and the incorporation of dU into DNA by polymerases. U lesions are repaired by base excision repair (BER) and BER initiated by uracil DNA glycosylase (UNG) is the predominant repair pathway for the removal of U incorporated into DNA by polymerases (4). UNG prefers a dA:T rich to dG:dC rich sequence, and single-stranded DNA (ssDNA) to double-stranded DNA (dsDNA), the latter, along with the very high turnover of UNG are consistent with a role in the rapid excision of U in replication forks (5). A second uracil DNA glycosylase, SMUG1, excises U from dU:dA pairs in dsDNA in nuclei derived from *ung-/-* knockout mice in the wake of the replisome and with slower kinetics. In short patch BER, abasic sites generated by UNG are hydrolyzed by apurinic/apyrimidinic endonuclease 1 (APE1), DNA polymerase β (Pol β) inserts a single T in the dsDNA template, and the nick is closed by Ligase III. In long patch BER, Pol β or DNA polymerase ε and the flap endonuclease FEN1 remove and replace several dNMPs in the dsDNA template (6).

ATR is an essential DNA damage response kinase activated at stalled and collapsed replication forks (7). ATR kinase signaling stabilizes stalled forks, reduces the rate of DNA replication, and increases nucleotide biosynthesis, thereby limiting the incorporation of dU by polymerases in cells treated with antimetabolites (8–10). ATR kinase inhibitors (ATRi) induce origin firing, inhibit deoxycytidine kinase, and induce the degradation of ribonucleotide reductase, thereby inhibiting nucleoside biosynthesis and nucleoside salvage, respectively, in otherwise unperturbed cells (11–13). ATRi therefore increase DNA synthesis and the ratio of dUTP:TTP and this causes deoxyuridine (dU) contamination. The ribonucleotide excision repair (RER) enzyme RNASE H2 is required for survival in cells treated with ATRi, which suggests ribonucleotides (rN) are also incorporated into DNA by polymerases in cells treated with ATRi (14,15). Since ATR kinase-dependent checkpoints are also inhibited, there is no signaling to limit DNA replication, and the incorporation of dU at DNA replication forks and ATRi are potent antimetabolites.

ATRi potentiate radiation-induced inflammation in patients and mouse models of cancer and induce the type 1 interferons (IFN-1) IFN-α/IFN-β, in cells (16–20). Structure-forming repeats are sites of replication fork collapse in cells treated with ATRi and this may impact both the structure and base composition of cytoplasmic DNA fragments induced by ATRi (19,21). While the base composition of these fragments may be molded by structure-forming repeats and the sequence preferences of UNG and SMUG1, the DNA damage that initiates ATRi-induced IFN-1 are not known. We hypothesized that dU and rN, concentrated in active replicons by ATRi-induced origin firing, may generate multiple damaged sites whose repair generates cytoplasmic nucleic acid fragments that induce IFN-1. Consistent with this premise, we recently showed that ATRi-induced dU incorporation and IFN-1 are reversed by low doses of thymidine, which implicates UNG- and SMUG1-initiated BER in the mechanism underlying ATRi-induced IFN-1 (10).

cGAS and RNA Polymerase III (Pol III) are pattern recognition receptors (PRRs) that identify dsDNA in the cytoplasm and initiate signaling pathways that induce IFN-1. cGAS recognizes B-form duplex DNA and catalyzes the production of cGAMP (cyclic GMP-AMP) which activates the adaptor protein STING (22–24). Activated STING recruits TANK binding kinase-1 (TBK1) which phosphorylates itself, STING and IRF3. IRF3, IRF7, and NF-κB induce IFN-β transcription. The major binding site for DNA in cGAS is a surface groove on the back side of the catalytic domain that is opposite to the substrate binding cleft (25,26). The interactions between cGAS and dsDNA are with the sugar-phosphate back bone of DNA and there is no known sequence specificty of the interaction. Pol III transcribes dsDNA in the cytoplasm to generate RNA that is recognized by RIG-1 (27). Once activated RIG-1 binds MAVS at the mitochondria and this results in TBK1 activation.

As a first step towards determining the molecular mechanism through which ATRi induce IFN-β *in vitro* and *in vivo*, we generated *ung-/-* knockout mouse cancer cell lines with the expectation that SMUG1 knockout would also be required to generate a phenotype. To our surprise, we discovered that UNG is preferentially required for ATRi-induced IFN-β, not because it is upstream of the endonuclease, but because cytoplasmic DNA fragments containing dU are not recognized by STING-dependent cytoplasmic DNA-sensing. Furthermore, *ung-/-* knockout sensitizes cells to the type II interferon, IFN-γ, *in vitro*, and generates responses to anti-PD-L1 in resistant tumors *in vivo*. These findings have important ramifications for cancer patients treated with antimetabolites and immunotherapy.

## RESULTS

### ATR kinase inhibitor-induced IFN-β is UNG-dependent

We generated *ung-/-* knockout mouse melanoma cell lines using CRISPR/Cas9. We selected melanoma cell lines as high UNG expression is associated with poor overall survival in melanoma **(Fig S1A)**. UNG1 and UNG2 have been considered mitochondrial and nuclear enzymes, respectively (6). However, recent work identified a distinct UNG1 isoform variant that is targeted to the nucleus and repairs genomic dU (28). This nuclear UNG1 variant, which in contrast to UNG2 lacks a PCNA-binding motif, may act on ssDNA through its ability to bind RPA. To avoid this complexity, we targeted both isoforms of UNG. Two clonal, homozygous *ung-/-* B16F10 cell lines (ΔUNG and ΔUNGΔPP) were used in this study. ΔUNG has a homozygous deletion that generates a frameshift and a stop codon; ΔUNGΔPP has a homozygous deletion that deletes two proline residues **(Fig S1B)**. ΔUNG and ΔUNGΔPP B16F10 have reduced glycosylase activity at dU:dA and dU:dG base pairs compared to WT, Cas9 expressing control cells **(Fig S2A)**.

First, we examined the sensitivity of WT and ΔUNG B16F10 to ATRi AZD6738 and ATM inhibitor AZD0156. ATRi-induced cell death was partially UNG-dependent in B16F10 **(Fig 1A; S3A,B)**. ATMi-induced cell death was also partially UNG-dependent **(Fig 1B)**. Next, we quantitated IFN-β induction in WT and ΔUNG B16F10 treated with 5 μM ATRi and 1 μM ATMi at 48 h, since these doses of inhibitors are equipotent with respect to cell viability. ATRi-induced IFN-β induction, and pTBK1 and pSTAT1 induction, were UNG-dependent in B16F10 **(Fig 1A; S3C-F)**. ATMi-induced IFN-β induction was not UNG-dependent **(Fig 1B)**. Thus, ATRi-induced, but not ATM-induced innate immune responses are UNG-dependent at doses of ATRi and ATMi that cause a ∼50% reduction in cell viability.

**Figure 1.**
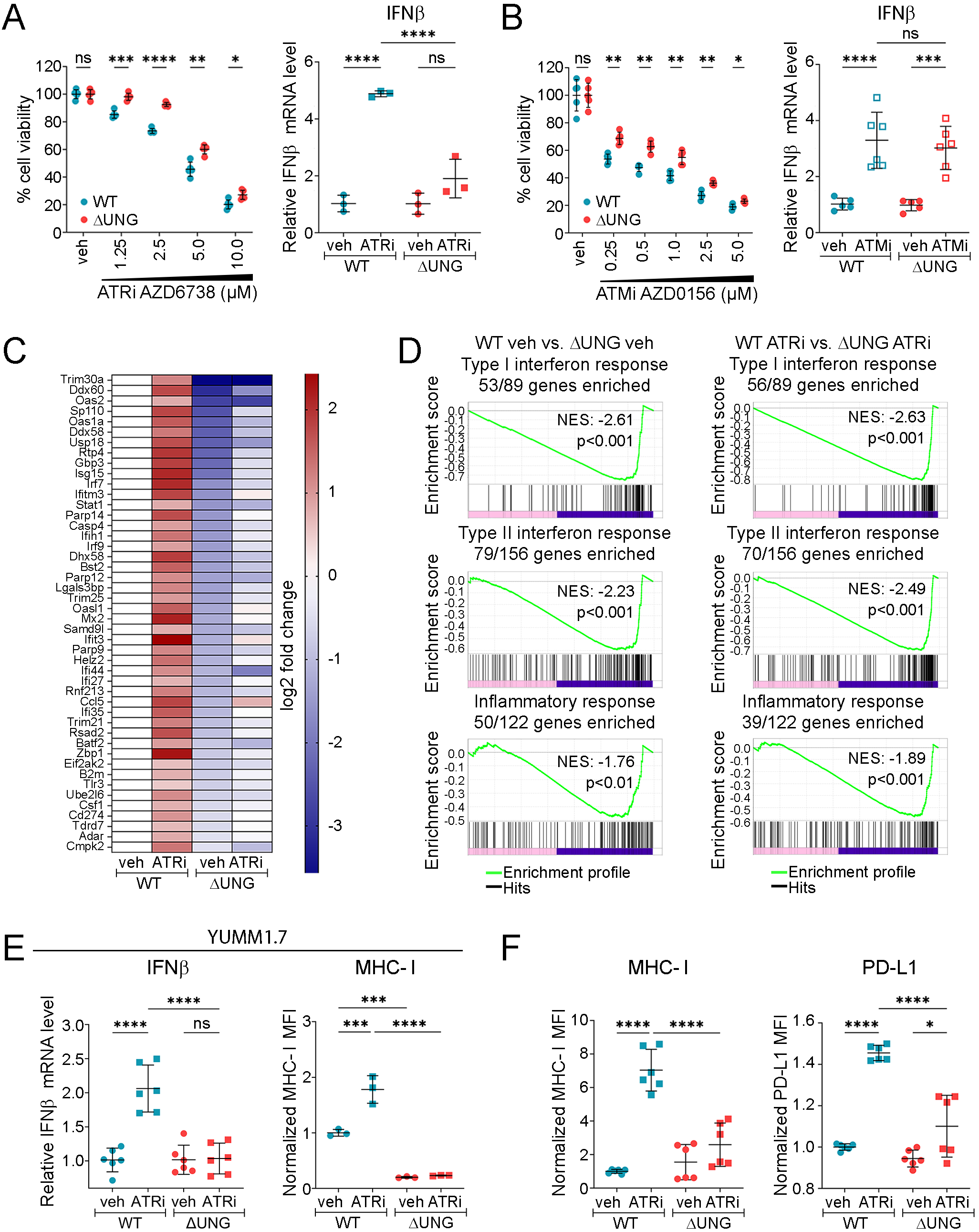
ATRi-induced innate immune signaling is UNG-dependent. **A,B.** WT and ΔUNG B16F10 were treated with ATRi AZD6738 **(A)** or ATMi AZD0156 **(B)**. Cell viability was determined using cell-titer glo at 72 h. Data were normalized to the mean of vehicle controls for each cell line and reported as % cell viability. Data from one representative experiment of two performed, each with four biological replicates. IFN-β mRNA expression was measured by qRT-PCR at 48 h with 5 µM ATRi or 1µM ATMi treatment. Expression was normalized to the mean of veh and are reported as the relative IFN-β mRNA level. Data from one representative experiment of two performed, each with three biological replicates. **C,D.** WT and ΔUNG B16F10 treated for 48 h with 5 µM ATRi. RNA sequencing was performed with two biological replicates. **C.** The inflammatory genes included in heatmap were the most significantly changed. **D.** Gene set enrichment analysis (GSEA) of the most significantly changed pathways. NES = normalized enrichment score. **E.** WT and ΔUNG YUMM1.7 treated with 5 µM ATRi for 16 h and IFN-β mRNA expression was measured by qRT-PCR. Data from two representative experiments, each with three biological replicates. Cell surface MHC-I expression was analyzed by flow cytometry. Median fluorescence intensities (MFI) were corrected for isotype control background and normalized to the mean of WT veh controls and are reported as normalized MFI. Data from one representative experiment of two performed, each with three biological replicates. **F.** WT and ΔUNG B16F10 were treated with 5 µM ATRi for 48h and cell surface MHC-I and PD-L1 expression were analyzed by flow cytometry. Normalized MFI data combined from two experimental replicates, each with three biological replicates. **A,B,E,F.** Data points represent individual biological replicates. Mean ± SD bars shown. *p<0.05, **p<0.01, ***p<0.001, ****p<0.0001, ns = not significant, determined by multiple unpaired t-tests (cell viability assay) or one-way ANOVA with Sidak’s multiple comparisons test (mRNA expression, flow cytometry).

We performed RNA sequencing to determine the genes altered by *ung* knockout and ATRi in an unbiased manner. RNA-seq revealed that the principal gene sets in which ΔUNG B16F10 cells are deficient are the type-I IFN response (53/89 genes at baseline), the type-II IFN response (79/156 genes at baseline), and the inflammatory response (52/122 genes at baseline) **(Fig 1C,D; S4).** The type-I IFN response (53/89 genes enriched), the type-II IFN response (79/156 genes enriched), and the inflammatory response were also deficient in ΔUNGΔPP B16F10 **(Fig S4G)**. UNG-dependent changes in ISG gene expression were validated using qRT-PCR **(Fig S3G,H)**.

ATRi-induced IFN-β induction was also UNG-dependent in YUMM1.7 **(Fig 1E)**, a second *Trp53* wild-type (WT) mouse melanoma cell line (29). Furthermore, ATRi-induced MHC-I induction was UNG-dependent in both YUMM1.7 and B16F10, and ATRi-induced PD-L1 induction was UNG-dependent in B16F10 cells **(Fig 1E,F)**. PD-L1 expression on YUMM1.7 was beneath the limit of detection and was therefore not included. In summary, UNG-deficient, *Trp53* WT mouse melanoma cells are deficient in type-I IFN, type II IFN, and inflammatory responses at baseline, and following treatment with ATRi.

### ATR kinase inhibitor-induced IFN-β is BER-dependent, but does not correlate with the concentration of dU in nuclear DNA

UNG-dependent IFN-β signaling in cells treated with ATRi may be a consequence of dU repair of ATRi-induced lesions that generates a PRR agonist and the accumulation of ATRi-induced damage in ΔUNG cells that generates a PRR antagonist.

The composition of the genome with respect to its constituent bases has not been quantitated and there are few assays that directly quantitate DNA lesions in the genome. We established the first quantitative assay for rA, rC, rG, rU, dA, dC, dG, dU, and T using a SCIEX6500+ LC-MS/MS system and developed a standard-of-practice to purify and digest DNA, free of RNA and R loops, into its constituent bases using enzymes and buffers that are free of dU contamination **(Fig 2A; S5; Supplemental Table 1)**. We were unable to detect rN in DNA above low levels of background contamination in the DNA Degradase enzyme. This is important as RNASE H2 knockout, which is expected to increase rN contamination in the genome, has been found to sensitize cells to ATRi in two unbiased screens (14,15).

**Figure 2.**
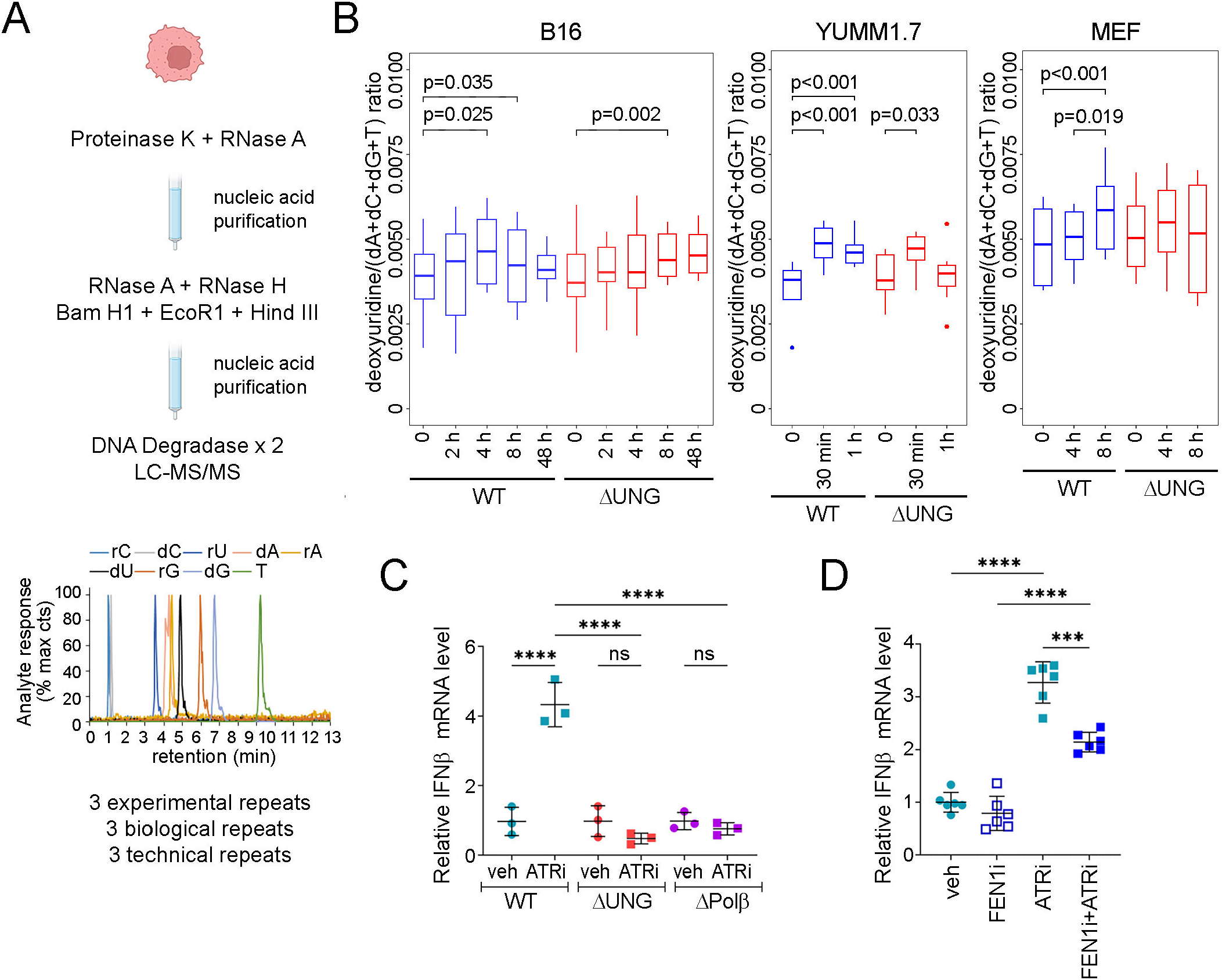
Steady state levels of dU in genomic DNA are similar in control and ΔUNG cells. **A.** Schematic representation of a novel LC-MS/MS assay for quantitation of ribonucleotide and deoxyuridine contamination in genomic DNA. **B.** Box plots show the ratio of dU: (dA + dC + dG + T) in genomic DNA purified from WT and ΔUNG B16F10, WT and ΔUNG YUMM1.7, and WT and ΔUNG MEF treated with 5 µM ATRi AZD6738 for indicated times. The data include 3 experimental replicates, each with 3 biological replicates per time/condition, and three technical replicates per biological replicate tested for pairwise differences in dU/dN ratio using a linear mixed model to account for experimental batch effects. P values are calculated for contrasts of interest. **C.** IFN-β mRNA expression was measured by qRT-PCR following 2.5 µM ATRi for 48 h or **(D)** 2.5 µM ATRi ± 5 µM FEN1i f for 24 h. Expression was normalized to the mean of veh controls and is reported as the relative IFN-β mRNA level. Data from one representative experiment of two performed, or two representative experiments combined, each with three biological replicates. **C,D.** Data points represent individual biological replicates. Mean ± SD bars shown. ***p<0.001, ****p<0.0001, ns = not significant, one-way ANOVA with Sidak’s multiple comparisons test.

The number of dU residues in genomic DNA in WT B16F10 was increased by ∼1,500 in 1×10^6^ bp after a 4 h treatment with 5 μM ATRi **(Fig 2B)**. The number of dU residues in genomic DNA in ΔUNG B16F10 cells increased by ∼820 in 1×10^6^ bp after a 4 h treatment with ATRi. The number of dU residues in genomic DNA in WT YUMM1.7 cells increased by ∼2,800 in 1×10^6^ bp after a 30 min treatment with 5 μM ATRi. The number of dU residues in genomic DNA in ΔUNG YUMM1.7 cells increased by ∼1,400 in 1×10^6^ bp after a 30 min treatment with ATRi. Thus, ATRi induced more dU in genomic DNA in WT than ΔUNG mouse melanoma cells.

We used MEF to quantitate dU in genomic DNA in cells with a full complement of DNA repair pathways. The number of dU residues in genomic DNA in WT MEF was increased by approximately 2,400 in 1×10^6^ bp after an 8 h treatment with 5 μM ATRi. The number of dU residues in genomic DNA in ΔUNG MEF was not increased after an 8 h treatment with 5 μM ATRi. Thus, ATRi induced more dU in genomic DNA in both WT than ΔUNG MEF, and WT than ΔUNG mouse melanoma cells.

We hypothesized that the ATRi-induced IFN-β signaling observed in WT, but not ΔUNG cells might be associated with futile cycles of long-patch BER. We treated WT, *ung-/-*, and *pol β-/-* MEF with 2.5 μM ATRi for 48 h and quantitated IFN-β mRNA expression. ATRi-induced IFN-β was UNG- and POLB-dependent **(Fig 2C; S2B)**. Next, we treated WT MEF with 2.5 μM ATRi and 5 μM FEN1i LNT 1 (30), an enzyme involved in long-patch BER, for 24 h. ATRi-induced IFN-β was, in part, FEN1-dependent **(Fig 2D)**. In contrast, and consistent with our data in mouse melanoma cells, ATMi-induced IFN-β was not UNG-dependent in MEF **(Fig S6A)**. Taken together these data show that ATRi-induced, but not ATMi-induced IFN-β induction is BER-dependent, and in part, long-patch BER-dependent in MEF.

### ATR kinase inhibitor-induced cytoplasmic DNA is increased in ΔUNG cells

The quantitation of dU in genomic DNA shows that steady state levels of dU in genomic DNA are similar in control and ΔUNG cells at baseline and following treatment with ATRi. This does not reconcile with the clear UNG-dependence of IFN-1 induction seen in mouse melanoma cell lines and MEF treated with ATRi **(Fig 1A,E,2C)**. The data above show that the UNG-dependent IFN-β response in cells treated with ATRi is associated with the repair of dU in genomic DNA by BER. This may generate a PRR agonist in WT cells. However, dU is not accumulating in genomic DNA in ΔUNG cells treated with ATRi compared to WT cells treated with ATRi, suggesting that dU in genomic DNA is repaired by a UNG-independent pathway in ΔUNG cells. We reasoned that a UNG-independent pathway may generate dU-rich DNA in the cytoplasm of ΔUNG cells and that dU-rich DNA may be a PRR antagonist. We therefore quantitated dU in cytoplasmic DNA in WT and ΔUNG cells treated with ATRi.

First, we treated WT and ΔUNG B16F10 with 5 μM ATRi for 48 h and enumerated micronuclei and micronuclei that stained positive for cGAS. There was no significant difference in the number of micronuclei or micronuclei containing cGAS in WT and ΔUNG B16F10 and ATRi increased both by similar amounts at 48 h (**Fig 3A-C**). Next, we purified and quantitated genomic DNA in the cytoplasm by qPCR. We observed a significant increase in genomic DNA in the cytoplasm of ΔUNG B16F10 treated with ATRi for 24 h, and an increase that was not significant in WT B16F10 treated with ATRi for 24 h (**Fig 3D,E**). Thus, while the number of micronuclei is similar in WT and ΔUNG B16F10 treated with ATRi, the amount of genomic DNA in the cytoplasm is modestly increased in ΔUNG B16F10 treated with ATRi than WT B16F10 treated with ATRi. This increase in genomic DNA in the cytoplasm of ΔUNG B16F10 treated with ATRi should increase rather than block the IFN-β response (compare Fig 1A,E to 3D,E).

**Figure 3.**
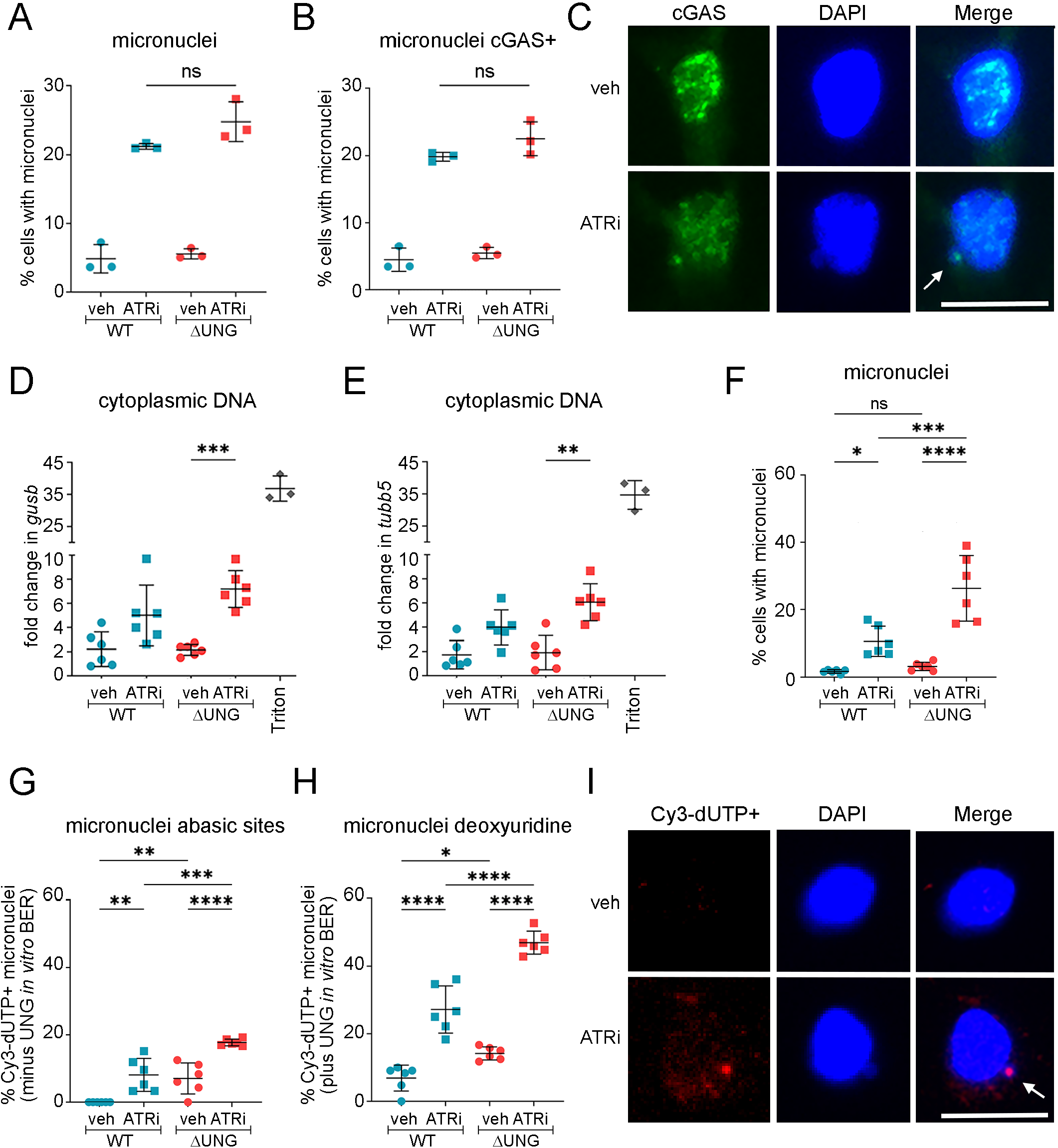
ATR kinase inhibitor-induced cytoplasmic DNA is increased in ΔUNG cells. **A-C.** WT and ΔUNG B16F10 were treated with 5 μM ATRi AZD6738 for 48 h and the percentage of cells that contain micronuclei and cGAS+ micronuclei was quantitated. A-B. Data points represent three biological replicates. **C.** Representative images; cGAS+ micronucleus denoted by the white arrow. Scale bar = 20 μm. **D,E.** WT and ΔUNG B16F10 were treated with 5 μM ATRi AZD6738 for 24 h and cytosolic DNA was isolated using digitonin fractionation. qPCR was performed to quantitate the relative change in the amount of the nuclear genes *gusb* and *tubb5* in the cytosolic fraction, normalized to whole cell DNA extracts. The triton fractionation control, in which the nuclear and mitochondrial membranes was permeabilized, is included in graphs but not in statistical testing. Data from one representative experiment of three performed, each with two biological replicates, and three technical replicates per biological replicate. **G-I.** WT and ΔUNG MEF were treated with 5 μM ATRi for 24 h. *In vitro* BER, with or without *E. coli* UNG, labelled abasic sites and dU sites in micronuclei with Cy3-dUTP (TTP analog). G. The percentage of total micronuclei following *in vitro* BER without UNG to idenfity abasic sites. H. The percentage of total micronuclei following *in vitro* BER with UNG to identify micronuclei containing deoxyuridine contamination. D-E. Data points represent a field of view with three chosen from each of two experimental replicates. I. Representative images; Cy3+ micronucleus denoted by the white arrow. Scale bar = 20 μm. **A,B,D,E,G-I.** Mean ± SD bars are shown. *p<0.05, **p<0.01, ***p<0.001, ****p<0.0001, ns = not significant, determined by one-way ANOVA with Sidak’s multiple comparisons test.

Since B16F10 is a pigmented cell line and melanin can obscure immunofluorescence analyses we quantitated dU in micronuclei in DNA in the cytoplasm of WT and ΔUNG MEF. First, we treated WT and ΔUNG MEF with 5 μM ATRi for 24 h and enumerated micronuclei. The number of micronuclei was significantly increased in ΔUNG MEF treated with ATRi compared to WT MEF treated with ATRi (**Fig 3F**). Next, we quantitated abasic sites in fixed MEF using *E. coli* DNA polymerase I. The number of abasic sites in micronuclei was significantly increased in ΔUNG MEF treated with ATRi compared to WT MEF treated with ATRi, but the difference was small (**Fig 3G**). Finally, we quantitated abasic sites and dU in micronuclei in fixed MEF using *E. coli* DNA polymerase I and *E. coli* uracil-DNA glycosylase (UNG). The number of abasic sites and dU in micronuclei was significantly increased in ΔUNG MEF treated with ATR compared to WT MEF treated with ATRi, and the difference was large (**Fig 3H,I**). In summary, micronuclei are increased in ΔUNG MEF treated with ATRi compared to WT MEF treated with ATRi and this should increase rather than block the IFN-β response (compare Fig 2C,D to 3F). However, more micronuclei in ΔUNG MEF treated with ATRi than WT MEF treated with ATRi contain dU-rich DNA. We therefore hypothesized that dU-rich DNA is a PRR antagonist.

### dU-rich cytoplasmic DNA antagonizes STING-dependent innate immune responses in cells

To test this hypothesis, we first investigated whether STING and Pol III innate immune responses are intact in WT and ΔUNG B16F10. The canonical STING agonist cGAMP and the canonical Pol III agonist poly(dA-T) induced IFN-β in both WT and ΔUNG B16F10 cells **(Fig 4A,B)**. Next, we generated STING knockout (ΔSTING) B16F10 cells using CRISPR/Cas9. We treated WT and ΔSTING B16F10 with 5 μM ATRi for 48 h **(Fig 4C; S6B)**. ATRi-induced IFN-β was largely STING-dependent, as previously observed (19,31).

**Figure 4.**
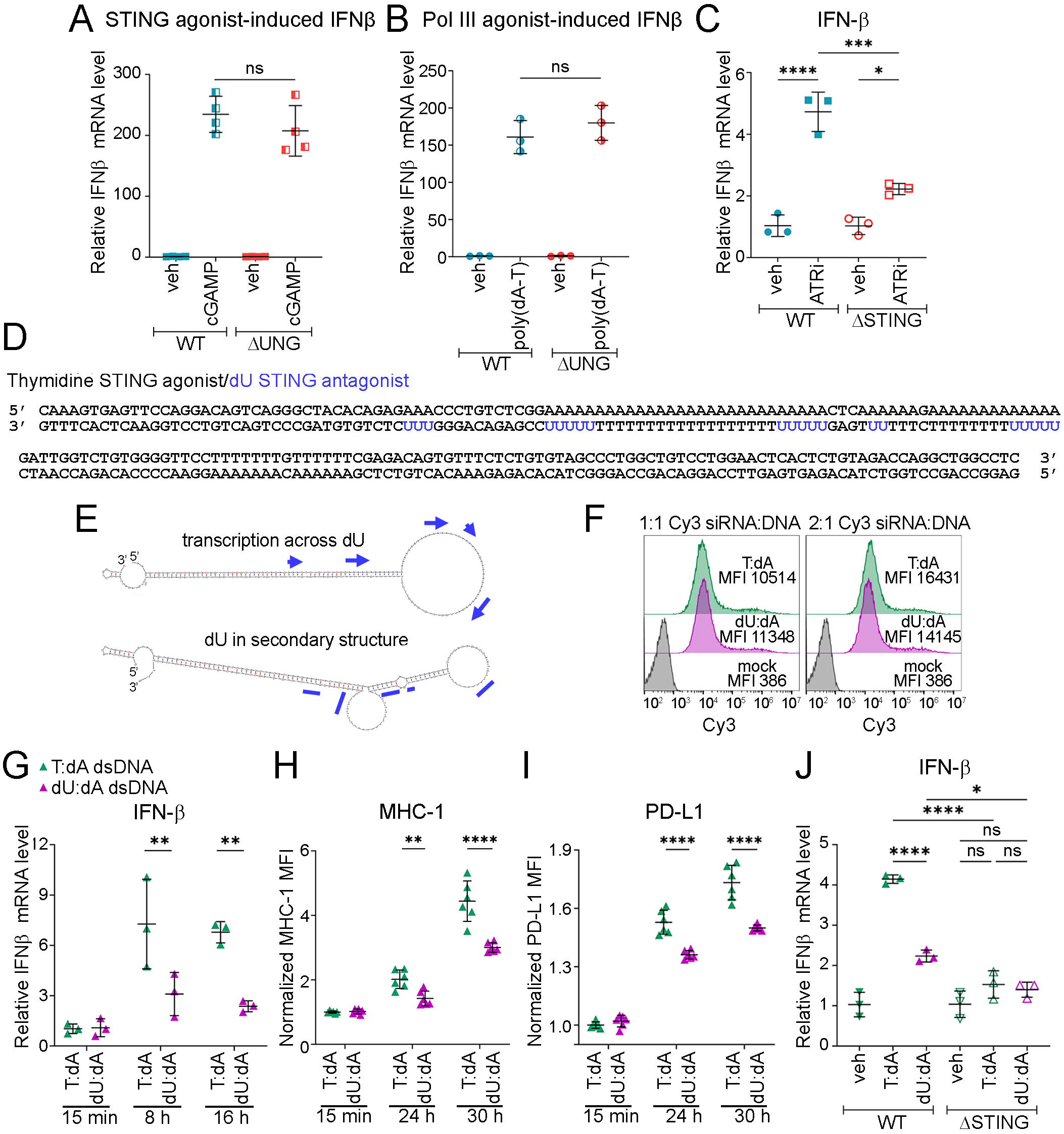
dU-rich cytoplasmic DNA antagonizes STING-dependent innate immune responses. **A,B.** Quantitation of relative IFN-β mRNA expression in WT and ΔUNG B16F10 treated with 10ug/ml cGAMP for 4 h or 0.2ug/ml poly dA:dT for 24 h. Data from one representative experiment of two performed, with four or three biological replicates. **C.** Quantitation of relative IFN-β mRNA level in WT and ΔSTING B16F10 treated with 5 μM ATRi AZD6738 for 48 h. Data from one representative experiment of two performed, each with three biological replicates. **D,E.** Sequence of dsDNA and secondary stucture of ssDNA. **F.** B16F10 were co-transfected with Cy3 siRNA and dsDNA oligos in 1:1 or 2:1 ratio and Cy3 MFI were determined using flow cytometry at 16 h post-transfection. Representative histograms with Cy3 MFI are shown. **G-I.** B16F10 were transfected with dsDNA oligos (0.1 µg/ml) for the indicated times. G. Quantitation of IFN-β mRNA level at 8 h and 16 h relative to the mean of 15 min conditions. Data from one independent experiment of two performed, each with three biological replicates. H-I. Cell surface MHC-I and PD-L1 expression were analyzed by flow cytometry at the indicated time points. MFI were corrected for isotype control background and normalized to the mean of the T:dA condition at the 15 min time point. H. Quantitation of normalized MHC-I MFI. I. Quantitation of normalized PD-L1 MFI. H-I. Data combined from two experimental replicates, each with three biological replicates, for a total of biological replicates. **J.** Control and ΔSTING B16F10 were transfected with dsDNA oligos (0.1 µg/ml) for 16 h. IFN-β mRNA levels relative to the mean the veh controls were quantified. Data from one representative experiment of two performed, each with three biological replicates. **A- C,G-J.** Data points represent biological replicates. Mean ± SD bars shown. A-C,J. *p<0.05, ***p<0.001, ****p<0.0001, ns = not significant, determined by one-way ANOVA with Sidak’s multiple comparisons test. G-I. **p<0.01, ****p<0.0001, ns = not significant, determined by two-way ANOVA with Tukey’s multiple comparisons test.

To directly determine whether dU impacts innate immune responses, we replaced thymidine bases with dU in synthetic 200 bp, double-stranded DNA (dsDNA) fragments previously shown to induce STING- and/or Pol III-dependent IFN-β in cells (19). The original sequence of the fragments was based on fragile sites expressed in cells treated with ATRi and these sequences were found to be associated with potential secondary structure (21). We selected one fragment that induced primarily STING-dependent IFN-β (original description: non-AT-rich#1 from mouse chromosome 4; 141468470-141468919) and one fragment that induced primarily Pol III-dependent IFN-β (original description: AT-rich#2 from human chromosome 9; 44235060-44235559), and for simplicity we refer to these fragments as STING agonist and Pol III agonist, respectively (19). In our design of the modified, dU-rich dsDNA fragments we were mindful of potential secondary structure and open reading frames for RNA polymerase III (Pol III), since Pol III synthesis of poly-U_>4_ is associated with pausing/termination (32).

The STING agonist **(Fig 4D)** has 23, 6, and 13 consecutive dA residues in the 5’ half of one strand and this is unlikely to be a template for Pol III. The complementary strand has 6 and 7 consecutive dA residues in the 5’ half of one strand and while this is also unlikely to be a template for Pol III, we choose to replace 20 T with dU on this strand 3’ to these consecutive dA residues as Pol III transcription on a template of pol-T_>4_ is possible. While we transfected dsDNA, we considered the predicted secondary structure of the single-stranded DNA in our design (13) **(Fig 4E).**

We transfected dA:T- and dA:dU-rich dsDNA STING agonists into WT B16F10 cells and quantitated transfection efficiency and innate immune responses. The transfection efficiency for dA:T- and dA:dU-rich dsDNA was similar **(Fig 4F)**. However, the relative induction of IFN-β, MHC-I, PD-L1, and ISGs (DDX58, Trim30A) was significantly reduced in cells transfected with dA:dU-rich dsDNA **(Fig 4G-I; S6C,D)**. At 16 h there was minimal induction of IFNβ by the dA:dU-rich dsDNA. The IFN-β induced by the dA:T-rich dsDNA, but not the dA:dU-rich dsDNA was STING-dependent **(Fig 4J)**.

Both strands of the Pol III agonist are potential templates for Pol III and we therefore replaced 40 T with dU in both strands **(Fig 5A)**. Again, we considered the predicted secondary structure of the single-stranded DNA in our design **(Fig 5B).** The transfection efficiency for dA:T- and dA:dU-rich dsDNA was again similar **(Fig 5C)**. Induction of IFN-β, MHC-I, PD-L1, and ISGs (DDX58, Trim30A) was again significantly attenuated by dA:dU-rich versus dA;T-rich dsDNA **(Fig 5D-F; S7A,B)**. However, at 16 h there was induction of IFN-β by the dA:dU-rich Pol III agonist dsDNA. The IFN-β and ISG (Trim30A) induced by the dA:T-rich dsDNA, but not the dA:dU-rich dsDNA was again STING-dependent **(Fig 5G; S7C)**. Furthermore, the MHC-I and PD-L1 induced by the dA:T-rich dsDNA, but not the dA:dU-rich dsDNA was STING-dependent **(Fig 5H)**.

**Figure 5.**
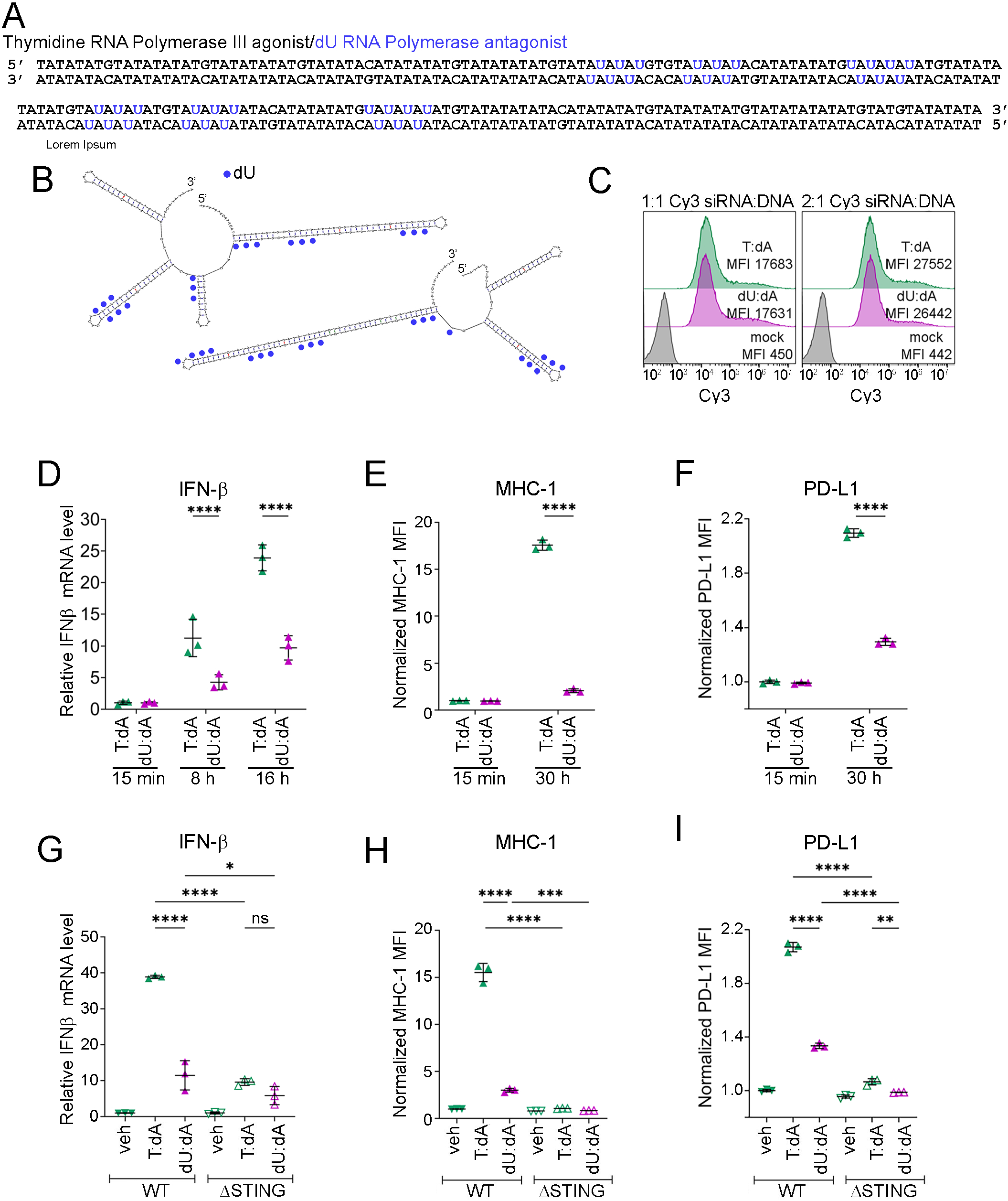
dU-rich, dA-T-rich cytoplasmic DNA antagonizes STING-dependent innate immune responses. **A,B.** Sequence of dsDNA and secondary stucture of ssDNA. **C.** B16F10 were co-transfected with Cy3 siRNA and dsDNA oligos in 1:1 or 2:1 ratio and Cy3 MFI were determined using flow cytometry at 16 h post-transfection. Representative histograms with Cy3 MFI are shown. **D-F.** B16F10 were transfected with dsDNA oligos (0.1 µg/ml) for the indicated times. D. Quantitation of IFN-β mRNA level at 8 h and 16 h relative to the mean of 15 min conditions. Data from one representative experiment of two performed, each with three biological replicates. E-F. Cell surface MHC-I and PD-L1 expression were analyzed by flow cytometry at the indicated time points. Raw MFI were normalized to the mean of the T:dA condition at the 15 min time point. E. Quantitation of normalized MHC-I MFI. F. Quantitation of normalized PD-L1 MFI. E-F. Data from one representative experiment of two performed, each with three biological replicates. **G-I.** Control and ΔSTING B16F10 were transfected with dsDNA oligos (0.1 µg/ml). G. IFN-β mRNA levels relative to the mean of veh controls were quantified at 16 h post-transfection. H-I. Cell surface MHC-I and PD-L1 expression were analyzed by flow cytometry at 30 h post-transfection. Raw MFI were normalized to the mean of the WT veh controls. H. Quantitation of normalized MHC-I MFI. I. Quantitation of normalized PD-L1 MFI. H-I. Data from one representative experiment of two performed, each with three biological replicates. **D-I.** Data points represent biological replicates. Mean ± SD bars shown. D-F. ****p<0.0001 determined by two-way ANOVA with Tukey’s (D) or Sidak’s (E-F) multiple comparisons test. G-I. *p<0.05, **p<0.01, ***p<0.001, ****p<0.0001, ns = not significant, determined by one-way ANOVA with Sidak’s multiple comparisons test.

### UNG knockout sensitizes cells to IFN-γ *in vivo*

Our *in vitro* data demonstrate that loss of UNG in B16F10 tumor cells disrupts tumor-intrinsic proinflammatory type I IFN signaling. We hypothesized that this would result in immunologically “cold” ΔUNG tumors *in vivo* that may grow more quickly. We confirmed that control and ΔUNG B16F10 cells and tumors grow at identical rates in vitro **(Fig S8A)** and in immunocompromised athymic nude mice **(Fig 6A; S8B)**. Surprisingly, ΔUNG tumors exhibited delayed, but more consistent tumor growth in immunocompetent C57BL/6 mice **(Fig 6A, B)**. This was evident at the earlier time point **(Fig 6B)**. While modest, these differences were significant and reproducible **(Fig 6A,B; S8C)**.

**Figure 6.**
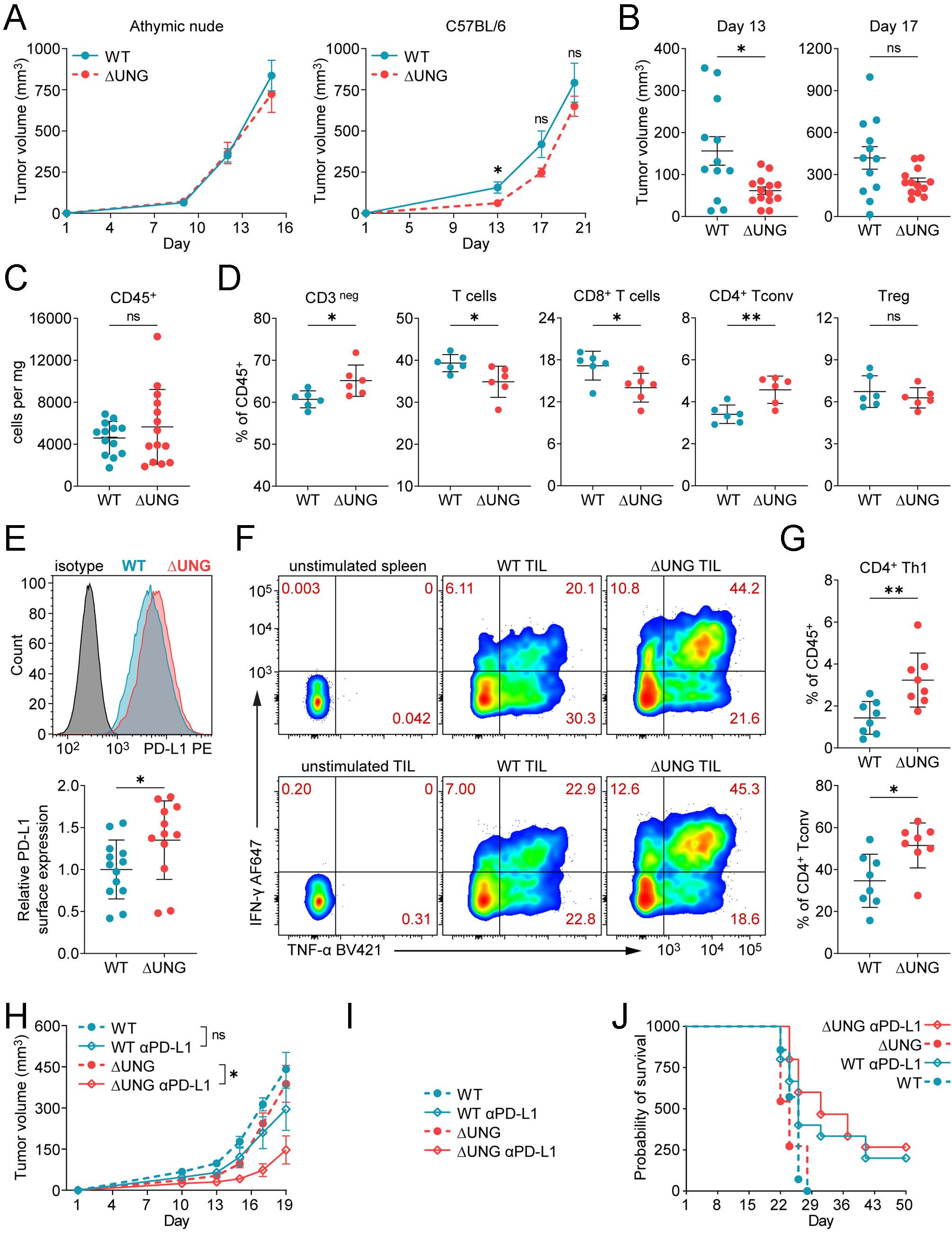
UNG loss alters the tumor-immune microenvironment and sensitizes B16 tumors to anti-PD-L1 therapy. **A.** Growth of WT and ΔUNG B16F10 tumors in immunocompromised athymic nude mice (left) or immunocompetent C57BL/6 mice (right). Mean tumor volumes and SEM bars shown. Data from one experiment per mouse strain. n = 19 athymic nude mice per group. n = 12 (WT) or 14 (ΔUNG) C57BL/6 mice. **B**. Individual WT and ΔUNG tumor volumes in C57BL/6 mice from day 13 and day 17 data in (A), with mean ± SEM bars shown. A-B. *p<0.05, **p<0.01, ns (not significant) by two-tailed, unpaired Welch’s t test at the indicated measurement time points. **C-D.** Flow cytometry immunoprofiling of WT and ΔUNG B16F10 tumors, grown in C57BL/6 mice, at day 14. C. Quantitation of the relative number of tumor-infiltrating CD45^+^ immune cells per mg of tumor stained. Data combined from 2 independent experiments, each with 6-8 mice per group, for total n = 13 (WT) or 14 (ΔUNG) mice. D. Quantitation of tumor infiltrating CD8^+^ T cells and conventional CD4^+^ T cells (Tconv) as percentages of the CD45^+^ immune cell infiltrate. Data from one experiment. n = 6 mice per group. **E.** Flow cytometry analysis of PD-L1 expression on control and ΔUNG B16F10 tumors (CD45neg cells) at day 14-15 in C57BL/6 mice. (Left) Histogram showing the median fluorescence intensity of PD-L1 PE staining on representative WT and ΔUNG B16F10 tumors, compared to isotype control staining. (Right) Quantitation of the relative cell surface expression of PD-L1 on WT and ΔUNG B16F10 tumors, normalized to the mean of WT control tumors. Data from 2 independent experiments, each with 4-8 mice per group, for total n = 13 (WT) or 12 (ΔUNG) mice. F-G. Flow cytometry analysis of IFN-γ and TNF-α production by CD4^+^ Tconv at day 15 following stimulation of control and ΔUNG B16F10 tumor infiltrates with PMA/ionomycin. **F.** Cytograms showing IFN-γ and TNF-α staining in representative stimulated control and ΔUNG B16F10 tumor infiltrates, compared to unstimulated control. **G.** Quantitation of tumor-infiltrating CD4^+^ Th1 cells, defined as IFN-γ^+^/IFN-γ^+^TNF-α^+^ CD4^+^ Tconv, as percentages of the CD45^+^ immune cell infiltrate (left) or parental CD4^+^ Tconv (right). Data from one experiment. n = 8 mice per group. C-E, G. Mean ± SD bars shown. *p<0.05, **p<0.01, ns (not significant) by two-tailed, unpaired t test. **H-I**. MHC-I cell surface expression was analyzed on B16 control and ΔUNG B16F10 treated in vitro with IFN-γ (0.1, 0.3, or 1.0 ng/mL) for 18 h. H. Histograms showing the median fluorescence intensity (MFI) of MHC-I PE staining on representative untreated or IFN-γ-treated (0.1 or 0.3 ng/mL) control and ΔUNG B16F10 samples, compared to isotype control staining. I. Quantitation MHC-I MFI, normalized to the mean of untreated WT cells. Mean ± SD bars shown. Data from one representative experiment with 3 biological replicates. **p<0.01, ****p<0.0001 by two-way ANOVA with Tukey’s multiple comparisons test. **J.** Control and ΔUNG B16F10 cells were injected (day 1) into C57BL/6 mice and mice were treated with 100 ug anti-PDL1 every 3 days for 6 doses, starting on day 2. Mean tumor volumes and SEM bars shown. *p<0.05, ns (not significant) by mixed-effects analysis of tumor growth data over time with Tukey’s multiple comparisons.

Next, we immunopurified the immune infiltrates in WT and ΔUNG tumors at day 14 after implantation. We profiled T cell populations in one experiment and profiled myeloid cell populations in a second experiment. Collectively, from the two experiments, we observed no significant difference in the total CD45^+^ immune cell infiltrate **(Fig 6C)**. However, we noted a significant decrease in CD8^+^ T cells and a significant increase in conventional CD4^+^ T cells (CD4^+^ Tconv) in ΔUNG tumors **(Fig 6D)**. Infiltration of regulatory T cells (Treg) was no different in WT and ΔUNG tumors **(Fig S8D)**. We also observed minimal differences in the myeloid populations profiled **(Fig S8E-G)**, except for reduced CD103^+^ dendritic cells (DC) in ΔUNG tumors **(Fig S8E)**. As CD103^+^ DC are important for priming the anti-tumor CD8^+^ T cell response (33), the reduction in this antigen-presenting population may explain the reduced CD8^+^ T cells in ΔUNG tumors.

We examined PD-L1 expression on CD45^neg^ tumor cells, as part of the myeloid cell profiling and an additional experiment. Contrary to our observation of reduced type I IFN-dependent expression of PD-L1 on ΔUNG cells in vitro, we noted increased tumor PD-L1 expression in ΔUNG compared to WT tumors in vivo **(Fig 6E)**. Since the type II-IFN, IFN-γ, is a primary mediator of PD-L1 upregulation by cells in vivo (34–36), we examined the cytokine competency of tumor-infiltrating T cells to determine whether increased T cell functionality is responsible for the elevated PD-L1 in ΔUNG tumors. We examined IFN-γ and TNF-α production by CD4^+^ Tconv and CD8^+^ T cells, and IL-17 production by CD4^+^ Tconv, following stimulation ex vivo with PMA/ionomycin **(Fig 6F,G; S8H,I)**. We saw no differences in IFN-γ/TNF-α-competent CD8^+^ T cells or IL-17-competent CD4^+^ Tconv (Th17 cells) in WT versus ΔUNG tumor-infiltrates **(Fig S8H,I)**. However, we observed significant increases in IFN-γ-competent (± TNF-α) CD4+ T cells, which represent the Th1 effector CD4^+^ T subset, as proportions of the CD45^+^ immune infiltrate and of the parent CD4^+^ Tconv population **(Fig 6G; S8I)**. Since ΔUNG tumors contain similar proportions of cytokine competent CD8^+^ T cells and fewer CD8^+^ T cells overall compared to WT tumors, increased CD4^+^ Th1 cells, rather than CD8^+^ T cells, are a likely source of IFN-γ driven PD-L1 expression in ΔUNG tumors.

Next, we aimed to determine whether control and ΔUNG B16F10 cells respond differently to IFN-γ. We treated cells *in vitro* for 18 hours with a dose range of IFN-γ (0.1, 0.3, and 1.0 ng/mL) and used upregulation of cell surface MHC-I and PD-L1 as readouts for responsiveness to IFN-γ **(Fig 6H-I; S8J,K)**. ΔUNG upregulated MHC-I significantly more than control B16F10 cells at the doses of IFN-γ tested **(Fig 6I; S8J)**. Upregulation of PD-L1 on WT and ΔUNG was similar at the two lower doses of IFN-γ but was increased on control cells treated 1.0 ng/mL IFN-γ compared to ΔUNG **(Fig S8K)**. These data support that ΔUNG cells are generally more sensitive to type II-IFN signaling despite exhibiting reduced type I-IFN signaling.

The B16F10 tumor model is known to be relatively resistant to immune checkpoint blockade, either via anti-PD-1 or anti-PD-L1 (37–39). Since, compared to the WT counterparts, ΔUNG tumors have altered tumor-immune microenvironments and ΔUNG cells exhibit increased sensitivity to IFN-γ, we questioned whether ΔUNG tumors have altered sensitivity to anti-PD-L1 therapy. We treated mice with anti-PD-L1 antibody every 3 days for 6 doses, beginning one day after implantation of WT or ΔUNG cells. While WT tumors treated with anti-PD-L1 exhibited some growth inhibition compared to WT controls, the differences were not statistically significant. Conversely, ΔUNG tumor growth was significantly inhibited by anti-PD-L1 therapy relative to growth of ΔUNG control tumors **(Fig 6J)**. Collectively, our data support that UNG activity in cancer cells is an important mediator of the tumor-immune microenvironment and of responsiveness to immunotherapy.

## DISCUSSION

Standard-of-care antimetabolites including 5-fluorouracil, pemetrexed, cytarabine, hydroxyurea, fludarabine, gemcitabine, methotrexate, 6-mercaptopurine, and capecitabine disrupt nucleotide pools and increase dU incorporation by DNA polymerases. We show that unrepaired dU in DNA blocks innate immune responses and potentiates responses to checkpoint inhibitors in mouse models of cancer. This would suggest that patients with low tumor UNG levels may respond to antimetabolites combined with checkpoint inhibitors, and patients with high tumor UNG levels may respond to UNG inhibitors

ATR kinase inhibitors (ATRi) induce origin firing, inhibit deoxycytidine kinase, and cause the degradation of ribonucleotide reductase and this disrupts both nucleoside salvage and nucleoside biosynthesis in undamaged cells (10,12). ATRi therefore increase DNA synthesis and decrease dNTP concentrations and thus dU contamination. Since ATRi inhibit ATR, there is no signaling to limit the incorporation of dU at DNA replication forks and ATRi are therefore potent antimetabolites (10).

UNG-dependent IFN-1 signaling in cells treated with ATRi could be a consequence of dU repair of ATRi-induced lesions that generates a PRR agonist, or the accumulation of ATRi-induced dU in ΔUNG cells that generates a PRR antagonist. Our previous observation that ATRi-induced IFN-1 is reversed by thymidine is consistent with the premise that dU at the replication fork is the initiating lesion (10).

UNG-initiated BER at dU concentrated across active replicons in cells treated with ATRi could generate extranuclear DNA recognized by PRRs. UNG-independent DNA repair could generate extranuclear DNA containing dU that is not recognized by PRRs. We suggest that both paradigms underlie our observations. First, ATRi induce origin firing and the unlimited incorporation of dU at replication forks, unlimited because ATR kinase-dependent cell cycle checkpoints are inhibited (10–12). Nevertheless, ATRi treatment is associated with the accumulation of ssDNA gaps at replication forks, presumably because ATRi-induced origin firing depletes dNTP pools causing polymerase stalling. Thus, in both WT and ΔUNG cells, ATRi induces nascent dsDNA regions that are separated by gaps at replisomes concentrated across active replicons. UNG co-localizes with PCNA at the replisome, UNG prefers ssDNA, dA-T-rich templates *in vitro*, and dU removal at the replication fork in nuclear extracts is UNG-dependent (4,40). While there are other DNA glycosylases that remove dU from genomic DNA, these do not appear to function at the replisome and the multiple damaged sites generated by ATRi may be refractory to the activity of alternate pathways. UNG cells may rapidly remove dU in WT generating nascent dsDNA regions containing abasic sites in WT cells separated by gaps at replisomes concentrated across active replicons. In contrast, dU is not removed in ΔUNG cells generating nascent dsDNA regions containing dU sites in ΔUNG cells separated by gaps at replisomes concentrated across active replicons.

Together our data show that ATM and ATRi induce cell death and DNA fragments that are recognized by PRRs in WT cells. But, in the case of ATRi, UNG must remove dU at the replisome to allow recognition by a PRR that activates STING. We show that ATRi-induced innate immune responses are STING-dependent in B16F10, a *Trp53* WT melanoma cell line. Furthermore, substitution of T:dA with dU:dA in DNA fragments generated by ATRi blocks STING-induced IFN-β, ISGs, MHC-1, and PD-L1. It has been shown previously that the T:dA STING agonist is cGAS-dependent. We therefore hypothesize that dU prevents the assembly of cGAS multimers in the cytoplasm, by disrupting a previously undescribed nucleotide sequence-dependent mechanism. We show here that cGAS localization in micronuclei and cGAMP-induced, STING-dependent innate immune responses are intact in ΔUNG cells. The nucleotide sequence-dependent mechanism that is disrupted by dU is an area of active investigation in our laboratory.

To determine whether this hypothesized mechanism has any consequences *in vivo*, we examined the growth of control and ΔUNG cells in immunoproficient mice. Remarkably, UNG knockout sensitized this checkpoint-resistant tumor model to anti-PD-L1. This was unexpected as increased innate immune responses and STING agonists are anticipated to potentiate checkpoint responses. Furthermore, our most dynamic UNG-dependent and thymidine-dependent variable is cell surface MHC-I expression which is dramatically reduced in ΔUNG cells and in agonists in which T:dA in substituted with dU:dA. We present data consistent with the hypothesis that dU:dA disrupts STING-dependent signaling and thereby sensitizes cells to IFN-γ. The responses to PD-L1 in a well-accepted checkpoint-resistant mouse model of cancer generated by UNG knockout are exciting and warrant further investigation given that antimetabolites are used to treat millions of cancer patients worldwide every year.

## METHODS DETAILS

### Cell Lines

Mouse embryonic fibroblasts (WT 92Tag, *pol β-/-* knockout 88Tag, *ung-/-* knockout 207Tag) were isolated from 14.5-day-old embryos and transformed by SV40 T-antigen expression (Dr. Robert W. Sobol) (41,42). MEFs, B16F10 (ATCC), and YUMM1.7 (Dr. Katherine Aird) were cultured in D-MEM, 10% fetal bovine serum, 1% Pen/Strep; YUMM1.7 media was supplemented with MEM Non-Essential Amino Acids. B16F10 and YUMM1.7 *ung-/-* knockout cells were generated using lentivirus expressing Cas9 and Cas9/sgRNA specific to mouse *ung*. Packaging vectors for the third-generation system pMDLg/pRRE, pRSV-Rev, and pMD2.G (all Addgene) and the shuttle vectors/transfer vectors were co-transfected into 293-FT cells using the TransIT-X2 Dynamic Delivery System (Cat# MIR 6003). Supernatant containing the lentivirus was collected after 48 hours (30). ΔSTING B16F10 were generated using gene knockout kit V2 (Synthego). A multi-guide specific to mouse *TMEM173* was transfected into cells according to the manufacturer’s instructions.

### Cell-survival analysis

Cells were seeded in 96-well microtiter plates and treated at 24 h. Cells were exposed to cell titer-Glo® 2.0 reagent for 10min at RT after the incubation interval indicated.

### RNA extraction and qPCR

Cells were harvested in Trizol and total RNA was extracted using the direct zol RNA kit (Zymo Research Corp). Total RNA was reverse transcribed using the Lunascript mastermix (NEB). qPCR was performed on a Quantstudio3 (Thermo).

### RNA-seq and data analysis

RNA was submitted to the Novogen Advancing Genomics facility (Sacramento, CA, USA) for RNA sequence analysis. Read alignment was performed using HISAT2 software using the mouse reference genome. FPKM was used to quantify the abundance of transcripts or genes. Differential gene expression analysis was performed using DEseq2 (for biological replicates) or edgeR (for no biological replicates) software with FDR correction by the Benjamini-Hochberg procedure with threshold [log2(foldchange)]>=1 & padj <=0.05. Group pathway analysis was performed using ClusterProfiler software for Gene Ontology analysis (Gene ontology analysis (http://www.geneontology.org/). Gene set enrichment analysis (GSEA 4.3.2) was utilized to rank based identification of most enriched pathway between groups. GO terms with padj <0.05 are significantly enriched, and the most significant terms were selected for display.

### Flow cytometry

Cells were blocked in normal mouse serum, stained with MHC-I or PD-L1 or isotype control antibodies, and efluor780 fixable viability dye. Cells were analyzed live or were fixed in 175 µL FluoroFix (BioLegend) prior to analyses. Single-stained OneComp eBeads (Invitrogen) were used as compensation controls for MHC-I and PD-L1. A single-stained mix of live and heat-killed (30 sec at 95°C) cells were used as the compensation control for efluor780. Acquisition was performed with a 4-laser CytoFLEX (Beckman Coulter), and analyses were performed in FlowJo V10.

For *in vivo* immune profiling experiments, WT or ΔUNG B16F10 tumors and spleens were harvested at the indicated time points. Splenocytes were used for single color controls, fluorescence-minus-one (FMO) controls, and for general gating. Tumor tissue (≤ 250 mg) was minced and then digested in Collagenase IV Cocktail containing Collagenase IV, DNase I, Soybean Trypsin Inhibitor. Tumor homogenate was smashed through a 70 μm cell strainer using the rubber plunger of a syringe. Spleens were mechanically dissociated between frosted glass slides and filtered through 70 μm cell strainers. Erythrocytes were lysed and cell suspensions were counted with a Scepter 3.0 (Millipore) and seeded at 1.2-2 x 10^6^ cells in 96-well round bottom plates for blocking and staining as follows: Fc receptors were blocked with anti-CD16/32 antibody, cells were stained with antibodies to surface antigens and eFlour780 viability dye or LIVE/DEAD fixable near-IR dead cell stain, samples were fixed and permeabilized in eBioscience Fixation/Permeabilization reagent (Invitrogen), and when performing nuclear (Ki67, Foxp3) or intracellular cytokine (IFN-γ, TNF-α, IL-17) staining, samples were stained with antibodies to nuclear/intracellular proteins. Brilliant Stain Buffer Plus (BD Biosciences) was added to antibody cocktails containing multiple Brilliant Violet dye conjugates to prevent polymer dye-dye interactions.

For measurement of cytokine-producing CD8^+^ and CD4^+^ T cells, prior to staining, cells were stimulated for 4 h with 1x eBioscience Cell Stimulation Cocktail (Invitrogen), which contains PMA and ionomycin, in complete D-MEM media. Unstimulated controls were treated with 1x eBioscience Cell Protein Transport Inhibitors cocktail (Invitrogen). Since PMA/ionomycin stimulation induces internalization of CD3 and the CD8/CD4 co-receptors, staining for CD3, CD4, CD8, and CD45 was performed post-fixation/permeabilization during intracellular cytokine staining. Uncompensated data were collected using a BD LSRFortessa 4-laser cytometer and BD FACSDiva software. Compensation and data analyses were performed in FlowJo V10 software. Single stained spleen samples with matching unstained cells or single stained OneComp eBeads (Invitrogen) were used for single color compensation controls. Fluorescence-minus-one (FMO) controls were used, where appropriate, to empirically determine gating. Gating strategies are shown **(Fig S9-S13)**.

### Quantification of nucleosides incorporated into the genome

Genomic DNA was prepared using the PureLink™ Genomic DNA Mini Kit (Invitrogen, K182002) protocol. 5 µg DNA was digested with RNase A (Thermo Scientific, EN0531), RNase H (New England BioLabs, M0297), Hind III, EcoRI, Bam HI. Digested DNA was purified using the GeneJET PCR Purification Kit (Thermo Scientific, K0702). DNA was eluted with RNase/DNase free water. Technical replicates consisting of 500 ng samples of DNA were digested into single nucleosides using DNA Degradase Plus (Zymo Research, E2021). Samples were heated to 96 °C. A second DNA Degradase Plus digestion was performed. To quantitate DNA constituent base composition, a fit-for-purpose LC-MS/MS assay was implemented on a 1290 Infinity II Autosampler and Binary Pump (Agilent) and a SCIEX 6500+ triple quadrupole mass spectrometer (SCIEX). Chromatographic separation was conducted on an Inertsil ODS-3 (3 µm x 100 mm 2.1 mm) reverse phase column (GL Sciences) at ambient temperature with a gradient mobile phase of methanol and water with 0.1% formic acid. MRM transitions of all analytes and isotopic internal standards were monitored to construct calibration curves. We were able to quantitate 1 rN per 20,000 bases from 1 mg DNA.

### Cytosolic DNA extraction and qPCR

Cells were resuspended in cytosolic extraction buffer containing 150 mM NaCl, 50 mM HEPES pH=7.5 (Corning), and 25 μg/mL digitonin (Cayman Chemical) and whole-cell homogenate was centrifuged for 15 min at 1000xg. Supernatant was recovered and centrifuged for 10 min at 20,000xg to recover 300 μL of cytosolic fraction. DNA was isolated using Quick-DNA Miniprep Plus Kit (Zymo Research). A sample containing 0.1% Triton-X100 in cytosolic extraction buffer (to permeabilize nuclear and mitochondrial membranes) was processed as described above and used as a positive control. DNA was amplified using the CFX Connect Real-time PCR system (Bio-Rad) and the PowerUp^TM^ SYBR^TM^ Green Master Mix (Applied Biosystems). The nuclear genes *Gusb* and *Tubb5* were amplified. The relative cytosolic fraction or Triton-X100 fraction DNA levels were normalized to whole-cell DNA levels using the delta-delta CT method.

### Micronuclei and cGAS staining

WT and ΔUNG B16F10 were treated and then fixed with 2 % PFA for 10 min at RT. Cells were blocked with 2% BSA and 0.1% Triton X-100 in PBS. Cells were incubated with anti-cGAS antibody overnight and secondary antibody for 1 h.

### In vitro BER for dU+ micronuclei

MEF were treated and then incubated with cytoskeletal buffer (100 mM NaCl, 300 mM sucrose, 10 mM PIPES pH 6.8, 3 mM MgCl_2_, and 0.5% Triton X-100) on ice for 5 min and fixed with 2% PFA for 10 min. MEF were permeabilized in 0.25 % Triton X-100. MEF were labeled with Cy3-dUTP using uracil DNA-Glycosylase, Endonuclease IV, Bst polymerase, Taq ligase, NAD+, 200nM dATP, 200nM dCTP, 200nM dGTP, and 200nM Cy3-dUTP.

### Innate immune agonists

Cells were treated with 10 ug/ml cGAMP in digitonin permeabilization solution for 10min and then incubated in complete media for 4 h, as described (43). Cells were treated with 0.2 µg/mL poly dA:dT complexed with lyovec^TM^ (Invivogen) for 24 h. dsDNA was synthesized by IDT and complexed with lyovec^TM^ (Invivogen). Cells were exposed to the complex in final concentration of 0.1 ug/ml in complete media.

### Animal experiments

Experiments were performed in accordance with protocols approved by the University of Pittsburgh Animal Care and Use Committee. Female 6-8 week old C57BL/6 mice and athymic nude mice were purchased from Jackson Laboratories. WT and ΔUNG B16F10 (2.5 x 10^5^ cells) in D-MEM were subcutaneously injected into the right hind flank of 8–10-week-old mice. Tumors were measured with digital calipers on the indicated days, and volumes calculated as volume = (length x width2)/2. For studies comparing the rates of growth of WT and ΔUNG B16F10 in nude mice or C57BL/6 mice, only tumors that established were included in measurements. For the study involving anti-PD-L1 therapy, mice were injected intraperitoneally with 100 µg anti-PD-L1 antibody (clone 10F.9G2, BioXCell inVivoPlus) diluted in 100 µL inVivoPure pH 6.5 Dilution Buffer (BioXCell), every 3 days for 6 doses, starting on day 2 after tumor cell injection on day 1. Control mice were injected intraperitoneally with 100 µL inVivoPure pH 6.5 Dilution Buffer. Since treatment began the day after tumor cell injection, all tumors, including those that did not take or that completely resolved, were included in the study group measurements. The tumor endpoint was reached when tumor volume exceeded 1000 mm^3^ or the tumor ulcerated. When multiple tumors within a group reached their endpoint, the entire study group was deemed to have reached the experimental endpoint. Mice that were euthanized due to tumor ulceration prior to the experimental endpoint were excluded from the reported tumor volumes.

### Statistics

The number of independent experiments and biological replicates (or mice), and the statistical tests performed (in GraphPad Prism 10, unless otherwise noted) are specified in figure legends. Experiments were performed at least 3 times unless otherwise noted in the text/legends. For data with multiple comparisons performed, stat bars and significance are only denoted for select comparisons of interest, but significance is adjusted for all comparisons made.

## Supporting information

Pandya Supplemental Table S1 4_3_2024

## ACKNOWLEDGEMENTS

This work was supported by CA236367 and CA204173 (CJB), CA240625 (KMA), CA148629, ES029518, ES028949, CA238061, and ES032522 (RWS) from the NIH. This project used the Cancer Pharmacokinetics and Pharmacodynamics Facility and the Cytometry Facility that are supported in part by award P30CA047904 from the NIH. Support was also provided by the Legoretta Cancer Center Endowment Fund (RWS). We thank Chris Koczor and Jianfeng Li (University of South Alabama) for discussions at the start of this project.

## AUTHOR CONTRIBUTIONS

PP, FPV, JEG, SP, AVC, YZ, designed and completed experiments in the Bakkenist lab. FPV and AVC completed all mouse experiments. JJD, RJ, completed the nucleoside quantitation in the Beumer lab. RB quantitated cytoplasmic DNA in the Aird lab. DI generated lentivirus, provided MEF, and performed the beacon assay (Supplemental Figure) in the Sobol lab. KMM and SMH provided essential, unpublished reagents and advice. ZH completed the biostatistics. All authors contributed to writing the paper.

