## Supplemental Figures and Extended Methods for "Deoxyuridine-rich cytoplasmic DNA antagonizes STING-dependent innate immune responses and sensitizes resistant tumors to anti-PD-L1 therapy"

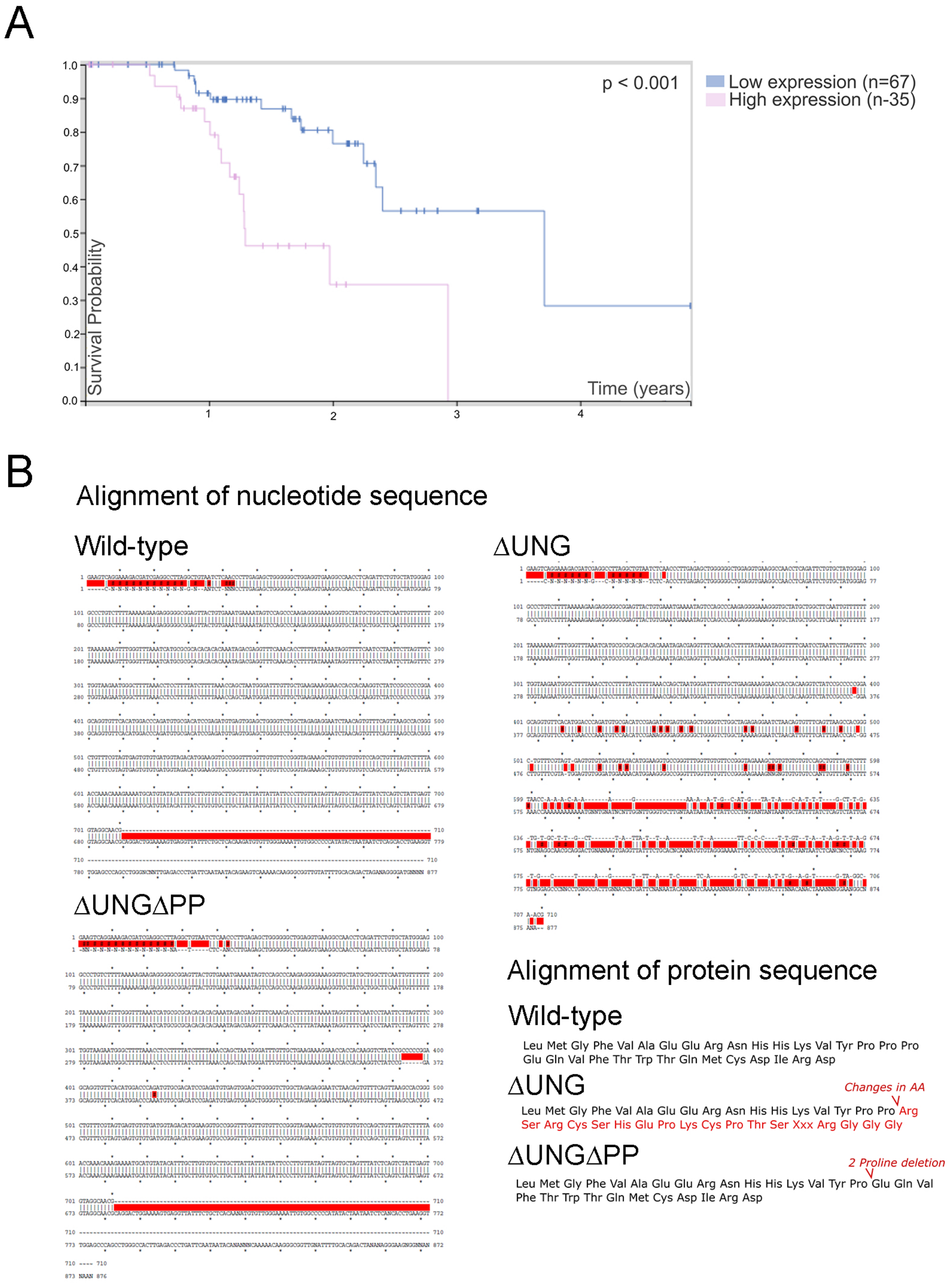


Pandya et al Figure S1

**Supplemental Figure S1A: UNG expression is a prognostic marker in melanoma.**

We selected melanoma cell lines as high UNG expression is associated with poor overall survival in melanoma **(Fig S1A)**. High vs. low UNG expression and survival probability are demonstrated in a Kaplan Meir plot from proteinatlas.org.

**Supplemental Figure S1B: Sanger sequence and amino-acid alignment for *ung-/-* B16F10 cell lines used in this study**.

Two clonal, homozygous *ung-/-* B16F10 cell lines (ΔUNG and ΔUNGΔPP) were used in this study. ΔUNG has a homozygous deletion that generates a frameshift and a stop codon; ΔUNGΔPP has a homozygous deletion that deletes two proline residues.

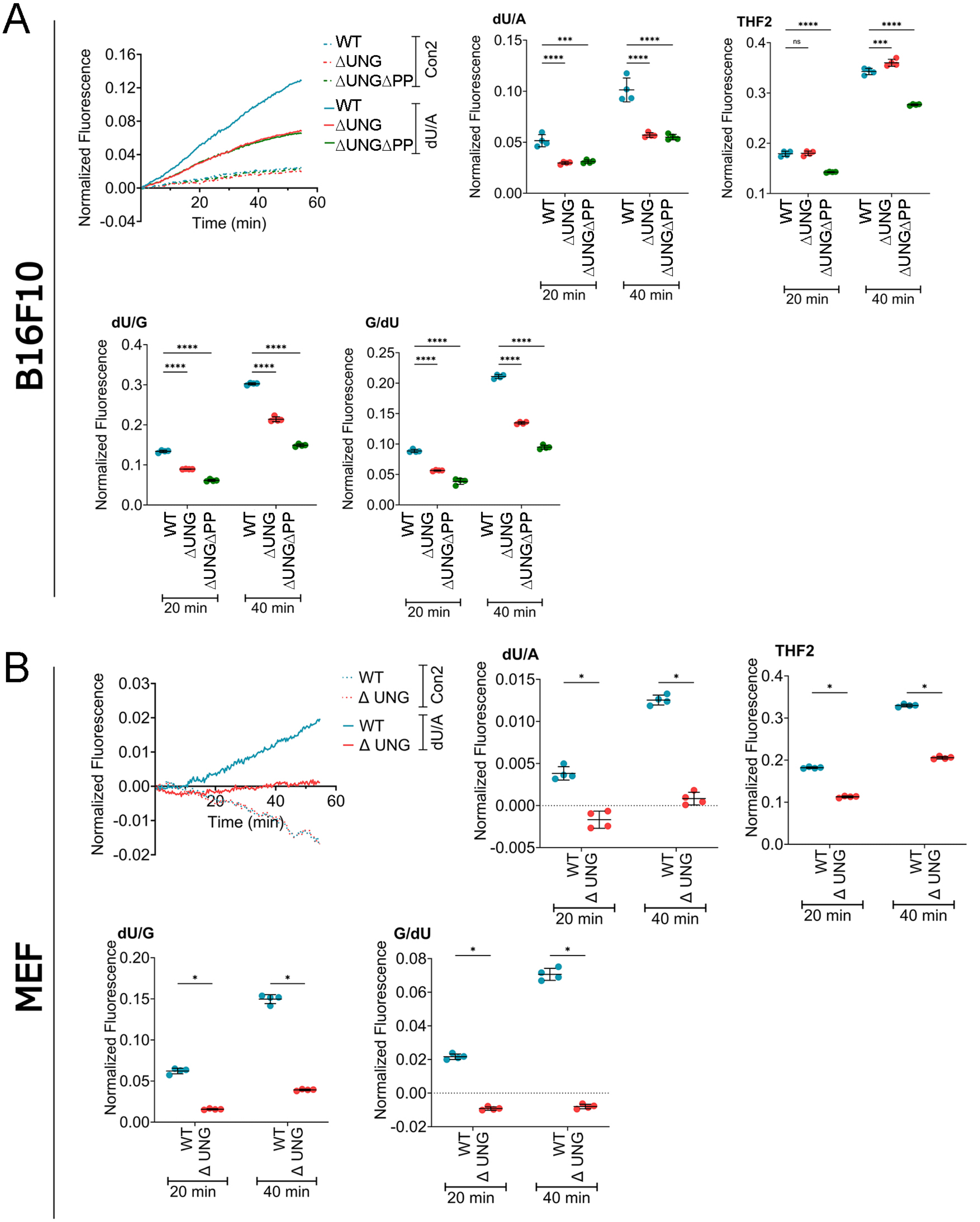


Pandya et al Figure S2

**Supplemental Figure S2A: ∆UNG and ∆UNG∆PP B16F10 have reduced glycosylase activity at dU:dA and dU:dG base pairs compared to WT, Cas9 expressing control cells.**

∆UNG B16F10 cells had significantly reduced activity against dU/dA, dU/dG, and dG/dU, as expected. ∆UNG B16F10 cells did not have reduced APE1 activity, as expected. ∆UNG B16F10 cells were used throughout this study.

∆UNG∆PP B16F10 cells had significantly reduced activity against dU/dA, dU/dG, and dG/dU, as expected. However, ∆UNG∆PP B16F10 cells had significantly reduced activity against THF, indicating that it also had reduced APE1 activity.

DNA repair beacons containing deoxyuridine or tetrahydrofuran (THF), mimicking an abasic sight and targeting APE1, are hairpins with a 6-Fam fluorophore on the 5' end and a Dabcyl non-fluorescent quencher on the 3' end. The dU/dA probe contains deoxyuridine opposite adenine. The dU/dG probe contains deoxyuridine opposite guanine. The dG/dU also contains a deoxyuridine opposite guanine, but in the reverse order. UNG activity induces fluorescence since it leads to the production of an AP site, subsequently cleaved by APE1.

**Supplemental Figure S2B: ∆UNG MEF have reduced glycosylase activity at dU:dA and dU:dG base pairs compared to WT MEF.**

∆UNG MEF had significantly reduced activity against dU/dA, dU/dG, and dG/dU, as expected. However, ∆UNG MEF had significantly reduced activity against THF, indicating that it also had reduced APE1 activity.

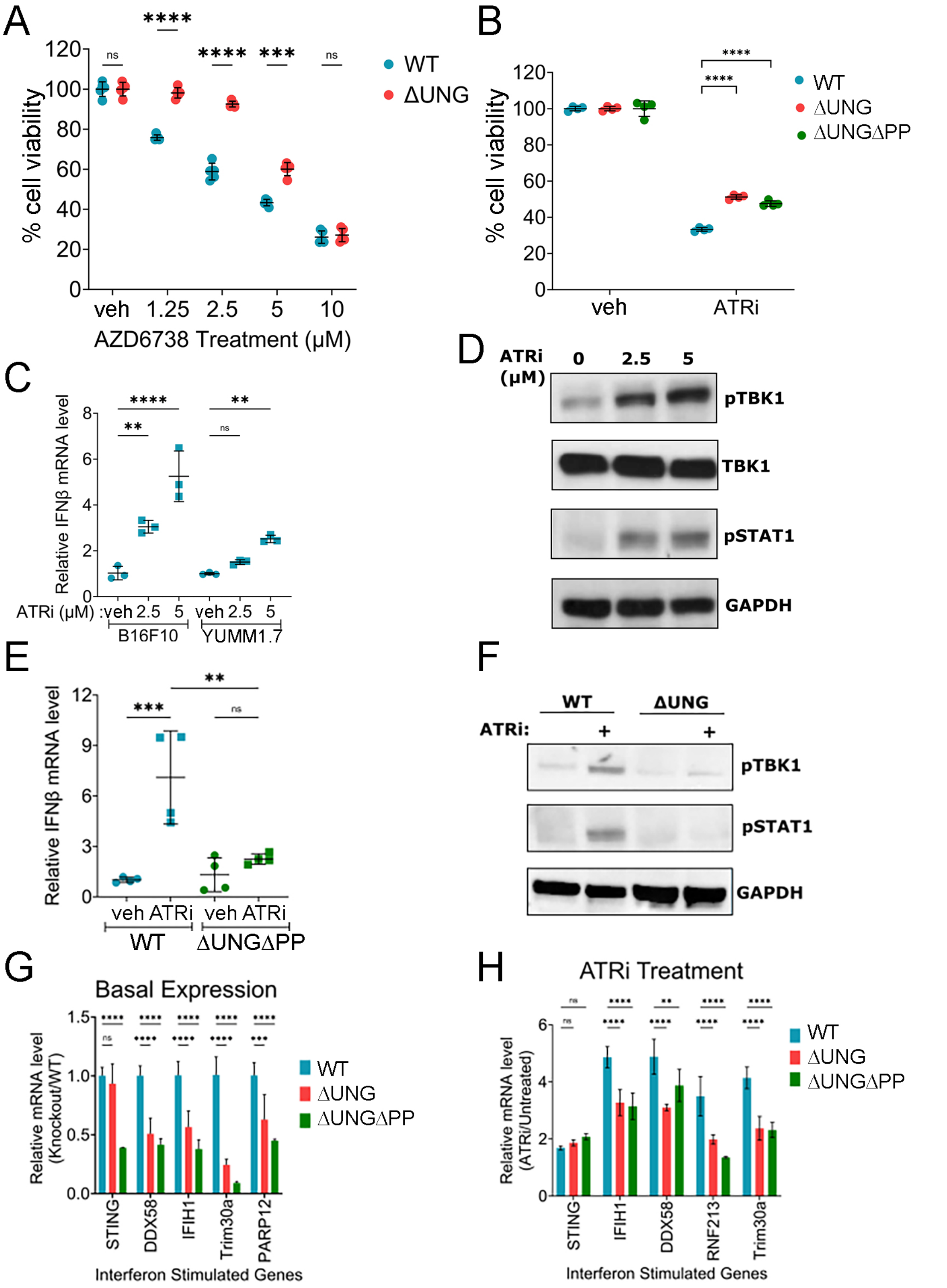


Pandya et al Figure S3

**Supplemental Figure S3: ATRi-induced IFN-β is reduced in ΔUNG and ΔUNGΔPP B16F10**

**(A-B)** WT and ΔUNG B16F10 (A) or WT and ΔUNGΔPP B16F10 (B) were treated with ATRi. Cell viability was determined using cell-titer glo at 48 h with the indicated dose of ATRi (A) or at 72h with 5 µM ATRi (B). Data were normalized to the mean of vehicle controls for each cell line and reported as % cell viability. Data from one experimental replicate (of two performed), each with 4 biological replicates. **(C)** B16F10 and YUMM1.7 cells were treated with 2.5 and 5 µM ATRi for 48h (B16F10) or 16h (YUMM1.7) and IFN-β was quantitated using qRT-PCR. Expression data were normalized to mean of veh for each cell line and represented as relative IFN-β mRNA level. Data from one experimental replicate (of two performed), each with 3 biological replicates. **(D)** WT B16F10 were treated with 2.5 and 5 µM ATRi for 48h and whole cell extracts were generated and immunoblotted for TBK1, pSTAT1, and GAPDH. **(E)** WT and ΔUNGΔPP B16F10 were treated with 5 µM ATRi AZD6738 for 48h and IFN-β was quantitated using qRT-PCR. Expression data were normalized to mean of veh for each cell line and represented as relative IFN-β mRNA level. Data combined from two experimental replicates, each with 2 biological replicates. **(F)** WT and ΔUNG B16F10 were treated with 5 µM ATRi for 48h and whole cell extracts were generated and immunoblotted for pTBK1, pSTAT1, and GAPDH. **(G,H)** WT, ΔUNG, and ΔUNGΔPP B16F10 were untreated (G) or treated with 5 µM ATRi (H) for 48h and ISGs were quantitated using qRT-PCR. Expression data were normalized to mean of veh and represented as relative ISG mRNA level. Data from one experimental replicate (of two performed), each with 3 biological replicates. (A-C,E,G,H) Mean ± SD bars are shown. *$p<0.05$, **$p<0.01$, ***$p<0.001$, ****$p<0.0001$, ns = not significant, determined by multiple unpaired t-tests (A); Two-way ANOVA with Dunnett's multiple comparison tests (B,G,H) or One-way ANOVA with Sidak's multiple comparisons tests (C,E).

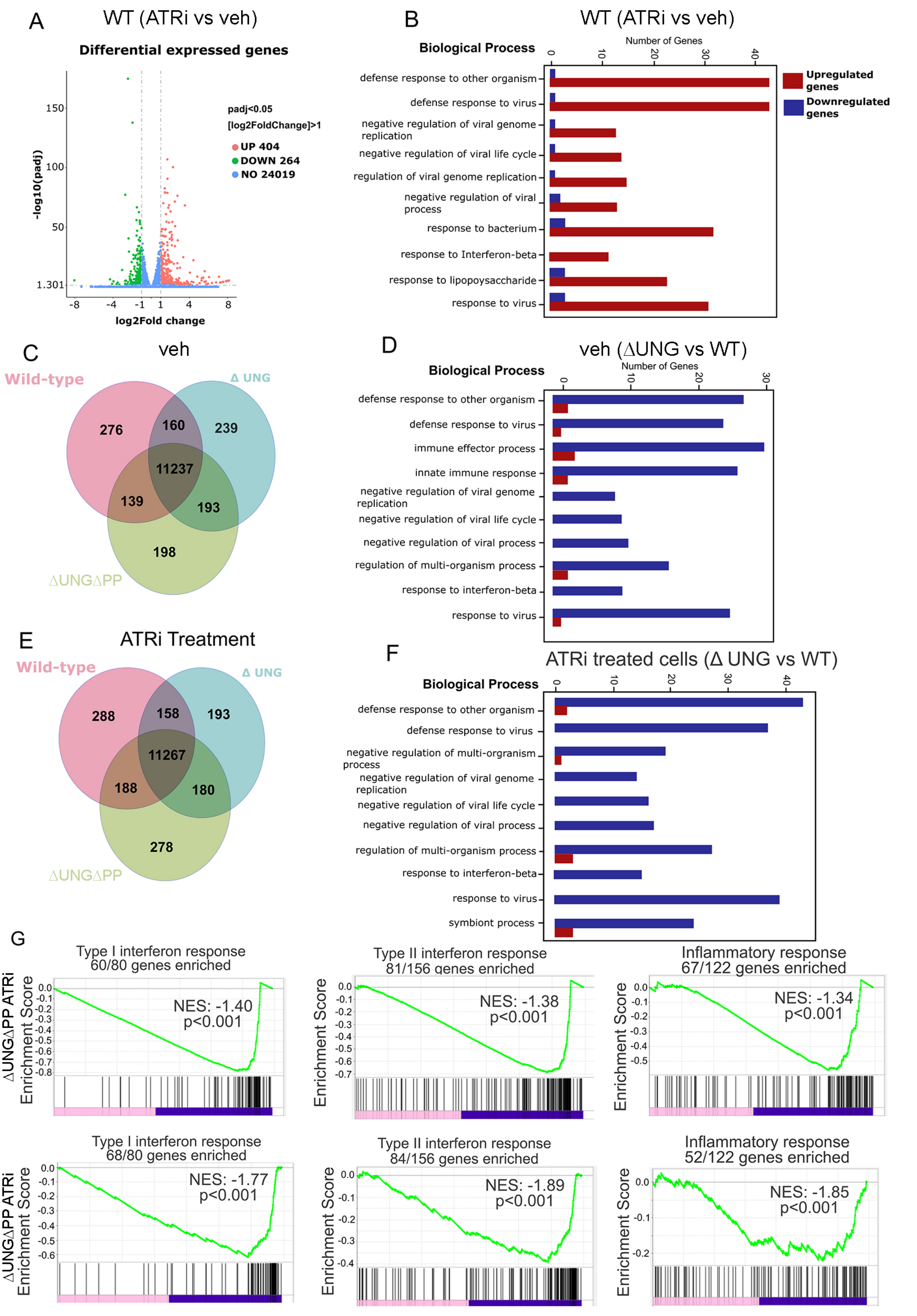


Pandya et al Figure S4

**Supplemental Figure S4: UNG-deficient B16F10 are deficient in type-I IFN, type II IFN, and inflammatory responses.**

B16F10 were treated with 5 μM ATRi for 48 h and RNA sequencing was performed with two biological replicates per condition. **(A)** Differentially expressed genes in veh vs. ATRi are shown in the volcano plot. Genes with [log2fold change]>1 and adjusted p-value <0.05 were considered significantly altered. Each point represents upregulated (red), downregulated (green), or unchanged (blue) genes. **(B, D, F)** Gene ontology analysis showing the top 10 most significantly altered biological processes in WT B16F10 treated with veh vs. ATRi, (B) WT vs. ΔUNG B16F10, (D) or ΔUNG B16F10 treated with veh vs. ATRi (F). **(C, E)** Venn diagrams of overlapping gene expression in WT, ΔUNG, and ΔUNGΔPP B16F10 treated with veh or ATRi. **(G)** Gene set enrichment analysis (GSEA) of the most significantly changed pathways in WT and ΔUNGΔPP B16F10 treated with veh or ATRi. NES = normalized enrichment score.

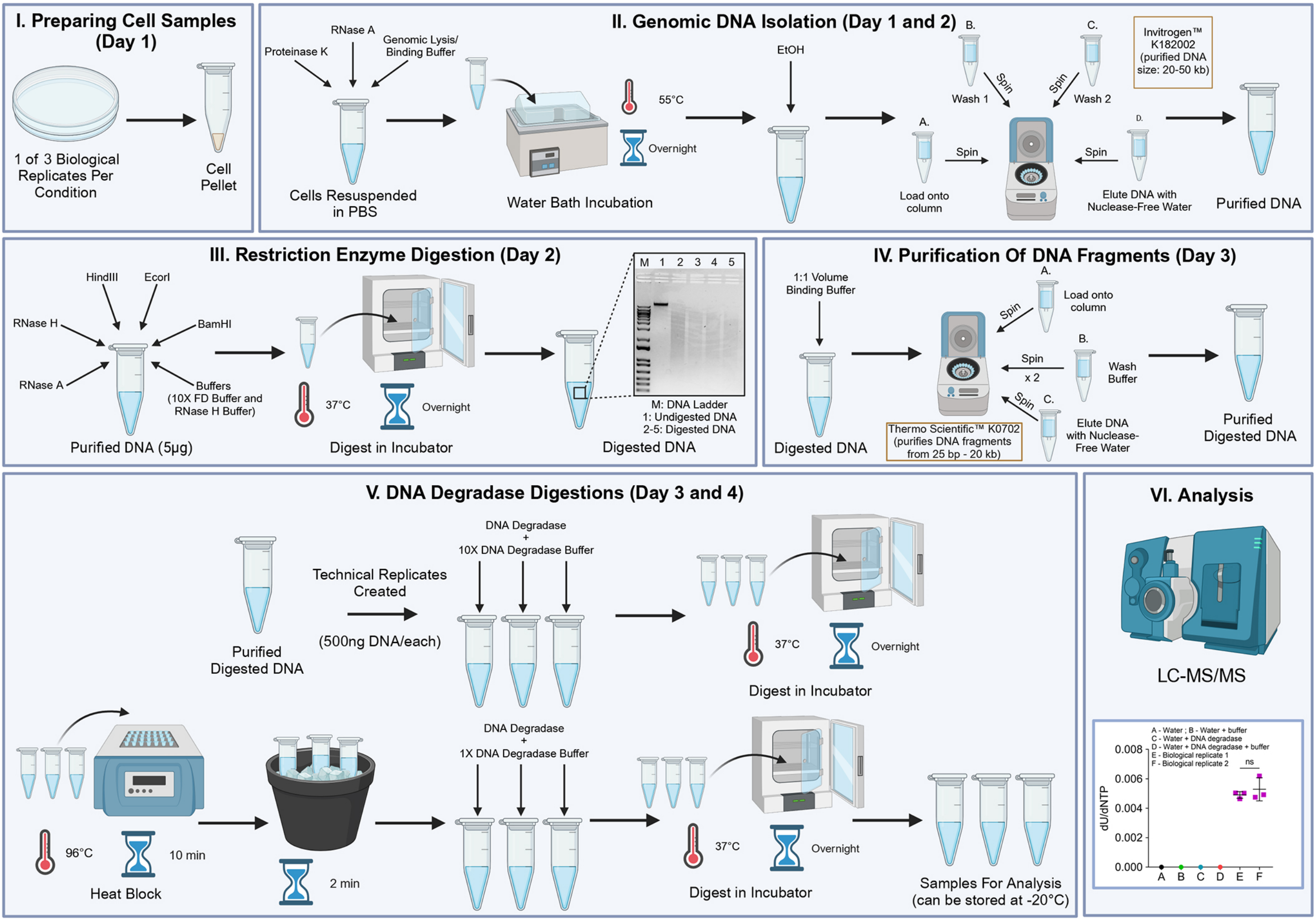


Pandya et al Figure S5

**Supplemental Figure S5: Preparation of samples to quantify nucleotides in the genome by LC-MS/MS.**

(I) Cells were harvested using trypsin and washed once with PBS-1X. (II) The cell pellet was resuspended in PBS-1X. An equal volume of DNA lysis buffer containing proteinase K and RNase A was added and the mixture was incubated at 55°C overnight. Genomic DNA was isolated using the standard protocol. (III) 5ug purified DNA was digested overnight at 37°C. (IV) DNA fragments were purified using a commercial kit. (V) DNA was quantitated using a NanoDrop. 500ng of each sample was digested with DNA degradase in triplicate overnight at 37°C. Samples were boiled. DNA degradase was added and the samples were digested overnight at 37°C. (VI) Samples were analyzed by LC-MS/MS. The bar graph shows one representative experiment with all controls.

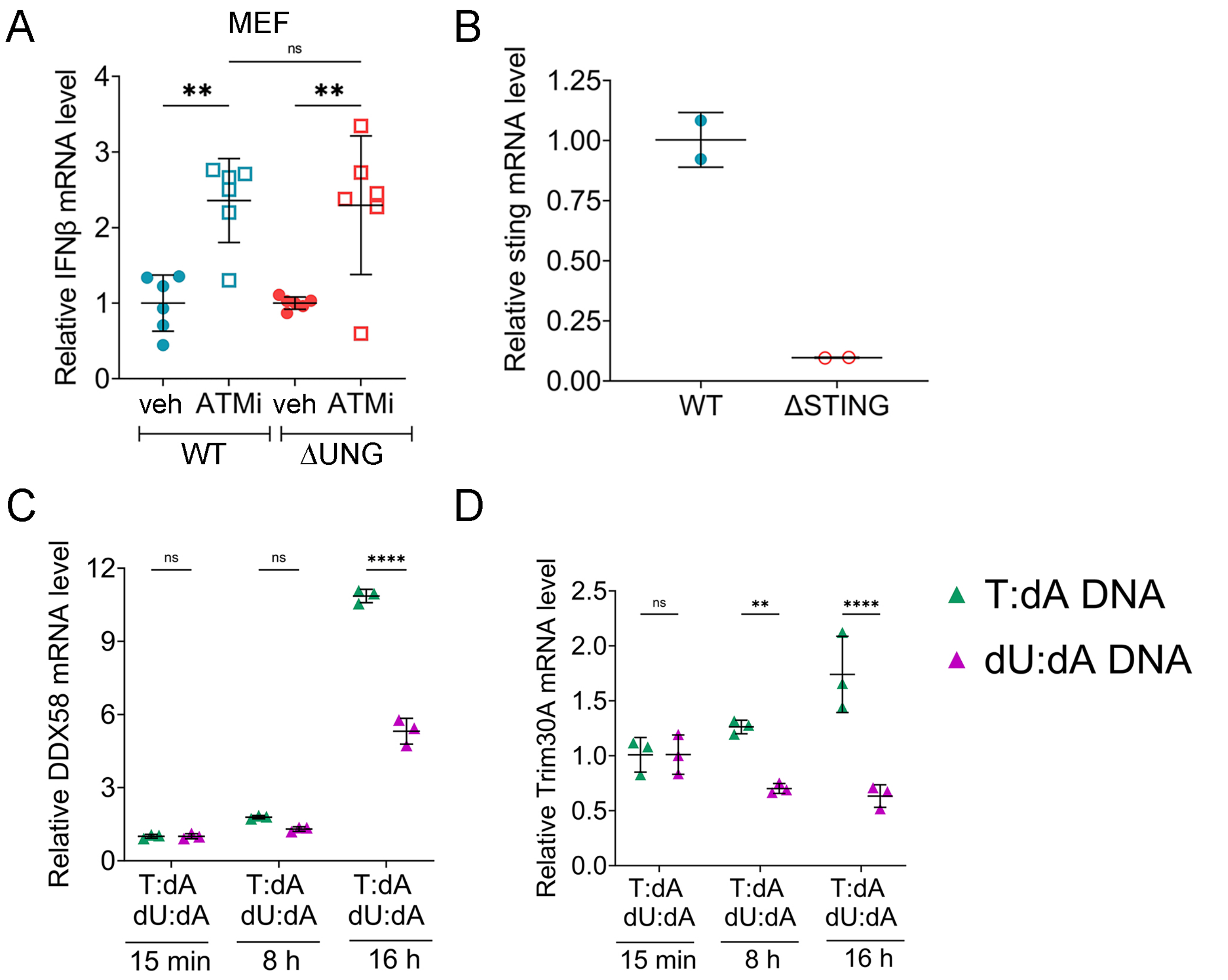


Pandya et al Figure S6

**Supplemental Figure S6: dU-rich cytoplasmic DNA antagonizes STING-dependent innate immune responses**

**(A)** WT and ΔUNG MEF were treated with 1 µM ATMi and IFN-β was quantitated using qRT-PCR. Expression data were normalized to mean of veh for each cell line and represented as relative IFN-β mRNA level. Data points are combined from two experimental replicates, each with 3 biological repeats. **(B)** STING mRNA expression was quantitated in WT and ΔSTING B16F10 using qRT-PCR. Relative change in STING mRNA expression from two biological replicates (n=2, mean ± S.D). **(C,D)** B16F10 were transfected with 0.1 µg/ml oligos (used in Fig.4) and *DDX58* and *Trim30A* mRNA expression were quantitated using qRT-PCR at 8 h and 16 h and normalized to the mean of 15 min conditions. Data from one experimental replicate (of two performed), each with 3 biological repeats. **p<0.01, ****p<0.0001, ns = not significant by (A) one-way ANOVA with Sidak’s multiple comparisons test or C,D) two-way ANOVA with Tukey’s multiple comparisons test.

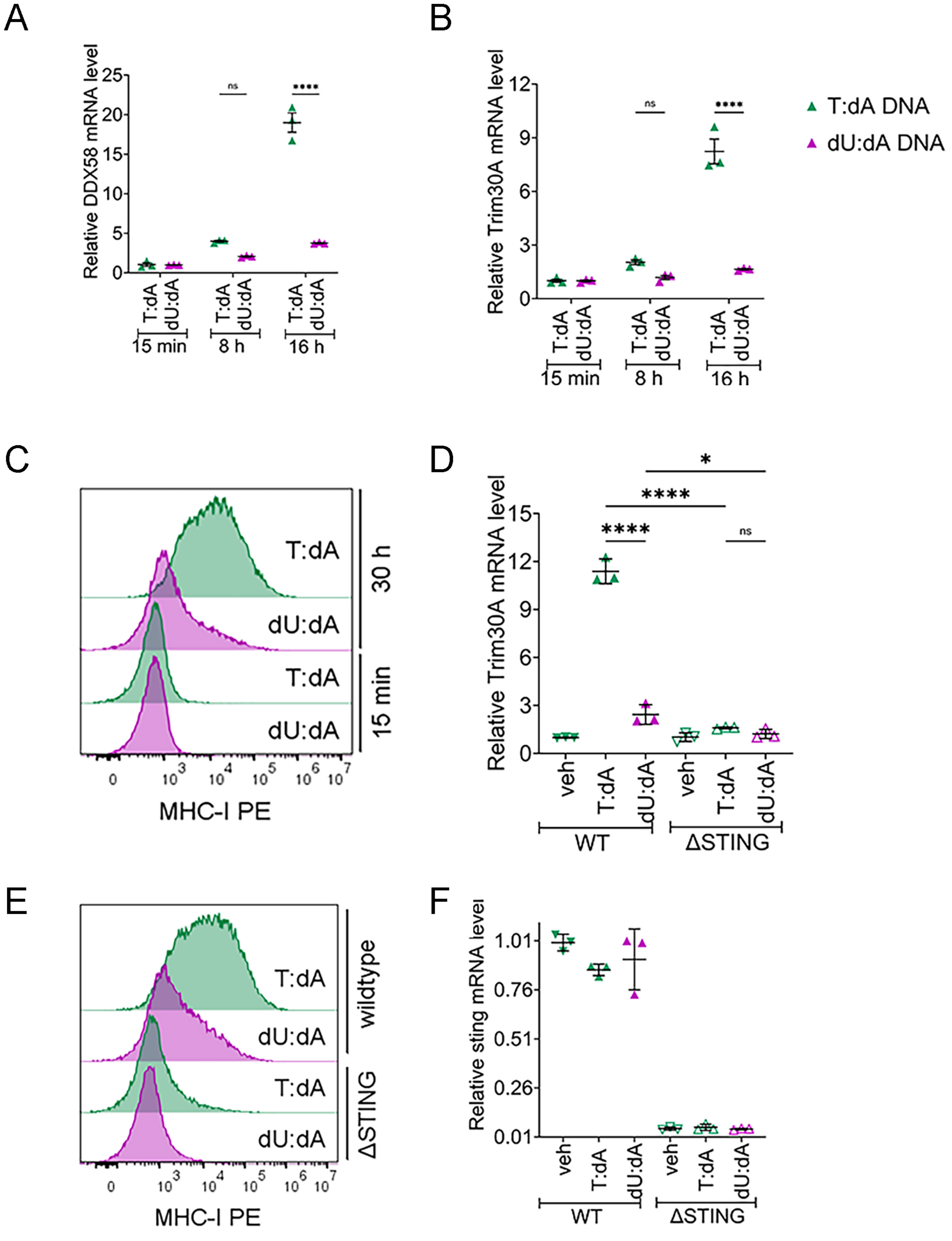


Pandya et al Figure S7

**Supplemental Figure S7:**

**(A,B)** B16F10 were transfected with 0.1 µg/ml oligos (used in Fig.5) and *DDX58* and *Trim30A* mRNA expression were quantitated using qRT-PCR at 8 h and 16 h and normalized to the mean of 15 min conditions. Data from one experimental replicate (of two performed), each with 3 biological replicates. **(C)** B16F10 were transfected with 0.1 µg/ml oligos (used in Fig.5). Representative histograms show MHC-I fluorescence intensity following 15 min and 30 h exposure to T:dA or dU:dA. **(D)** WT and ∆STING B16F10 were transfected with 0.1 µg/ml oligos (used in Fig.5) and *Trim30A* expression was measured and quantitated relative to the mean of WT veh controls. Data from one experimental replicate (of two performed), each with 3 biological repeats. **(E)** WT and ∆STING B16F10 were transfected with 0.1 µg/ml oligos (used in Fig.5). Representative histograms show MHC-I fluorescence intensity following 30 h exposure to T:dA or dU: dA oligo. **(F)** WT and ∆STING B16F10 were transfected with 0.1 µg/ml oligos (used in Fig.5). STING mRNA expression was quantitated using qRT-PCR relative to the mean of WT veh controls. Data from one experimental replicate (of two performed), each with 3 biological repeats. *$p<0.05$, ****$p<0.0001$, ns = not significant by (A,B) two-way ANOVA with Tukey's multiple comparisons test or (D,F) one-way ANOVA with Sidak's multiple comparisons test.

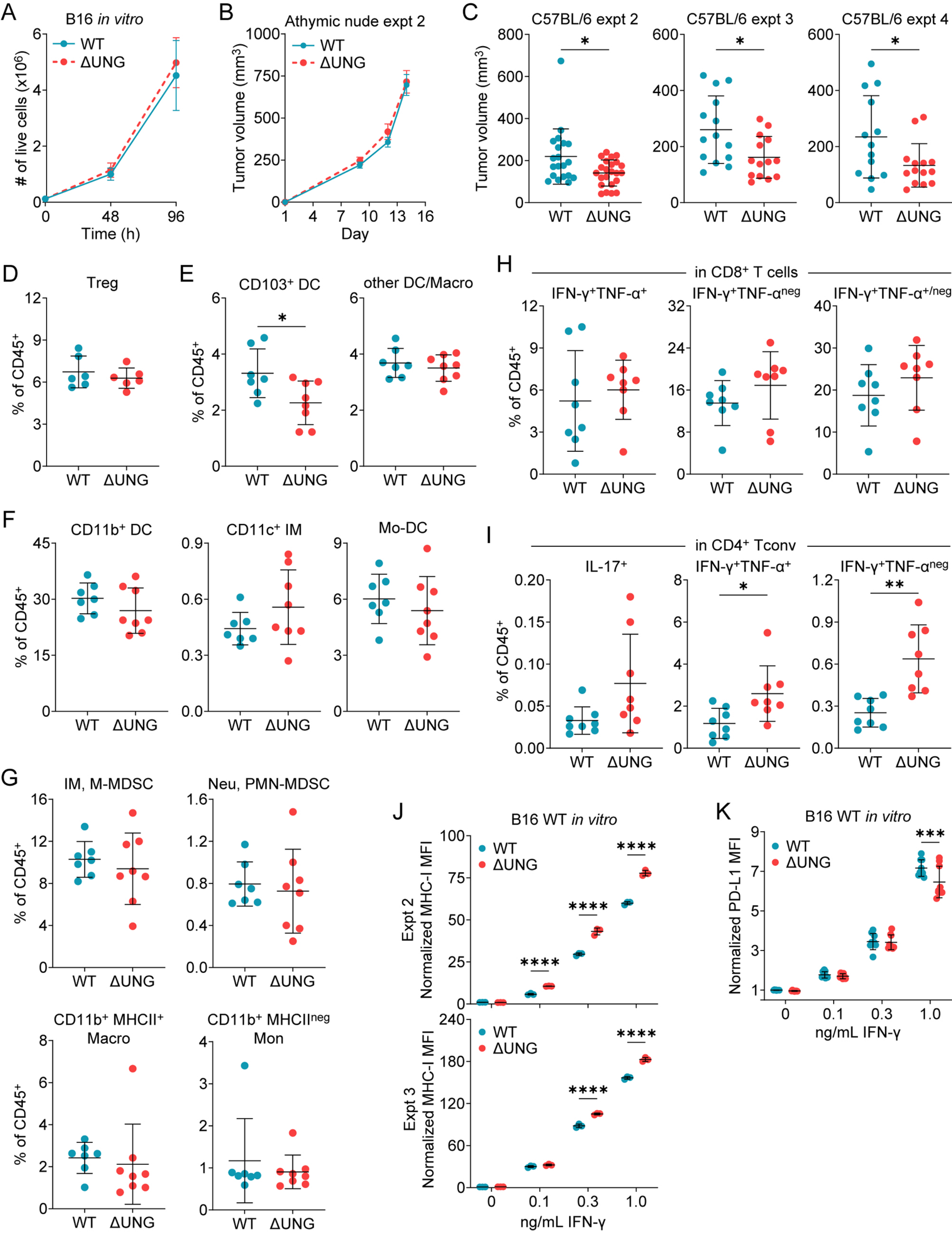


Pandya et al Figure S8

**Supplemental Figure S8. UNG loss alters the B16 tumor-immune microenvironment.**

**(A)** *In vitro* growth of WT and ΔUNG B16F10 cell lines. Cells were seeded at equal density and counted at 48 and 96 hours. **(B)** Repeat experiment for growth of WT and ΔUNG B16F10 tumors in immunocompromised athymic nude mice. Mean tumor volume ± SEM bars shown. n = 18 (WT) or 19 (ΔUNG) mice. **(C)** Independent repeat experiments for growth of WT and ΔUNG B16F10 tumors in immunocompetent C57BL/6 mice. Shown are individual WT and ΔUNG tumor volumes day 13 from three independent repeat experiments, with mean ± SD bars shown. n = 13-20 (WT) or 14-23 (ΔUNG) mice per experiment. *$p<0.05$ by two-tailed, unpaired Welch's t test. (**D-I)** Flow cytometry immunoprofiling of WT and ΔUNG B16F10 tumors, grown in C57BL/6 mice, at day 14. **(D)** Quantitation of tumor infiltrating regulatory T cells (Treg) as percentages of the $CD45^+$ immune cell infiltrate. Data from one experiment. n = 6 mice per group. **(E)** Quantitation of $CD103^+$ dendritic cells (DC) and other dendritic cells or macrophages (other DC/macro, $CD11c^+CD103^{neg}CD11b^{neg}$) within the $CD11c^+CD11b^{neg}$ myeloid population. **(F)** Quantitation of myeloid subsets within the $CD11c^+CD11b^+CD103^{neg}$ population based on expression of Ly-6C and MHC-II. The subsets include: $CD11c^+CD11b^+$Ly-$6C^{neg}$MHC-$II^{neg}$ cells defined as $CD11b^+$ dendritic cells ($CD11b^+$ DC); $CD11c^+CD11b^+$Ly-$6C^+$MHC-$II^{neg}$ cells defined as $CD11c^+$ inflammatory monocytes ($CD11c^+$ IM); and $CD11c^+CD11b^+$Ly-$6C^+$MHC-$II^+$ cells defined as monocyte-derived dendritic cells (Mo-DC). **(G)** Quantitation of myeloid subsets within the $CD11b^+CD11c^{neg}$ population based on expression of Ly-6C, Ly-6G, and MHC-II. The subsets include: $CD11b^+CD11c^{neg}$Ly-$6C^{hi}$Ly-$6G^{neg}$ inflammatory monocytes (IM) or monocytic (mononuclear) myeloid-derived suppressor cells (M-MDSC); $CD11b^+CD11c^{neg}$Ly-$6C^{int}$Ly-$6G^+$ neutrophils (Neu) or granulocytic/polymorphonuclear myeloid-derived suppressor cells (PMN-MDSC); $CD11b^+$MHC-$II^{neg}$ (and CD11c/Ly-6C/Ly-6G neg) monocytes ($CD11b^+$MHC-$II^{neg}$ Mon); and $CD11b^+$MHC-$II^+$ (and CD11c/Ly-6C/Ly-6G negative) macrophages ($CD11b^+$MHC-$II^+$ Macro). **(E-G)** Data from one experiment with n = 7-8 mice per group. **(H-I)** Flow cytometry analysis of IFN-γ and TNF-α production by $CD8^+$ T cells or IFN-γ, TNF-α, or IL-17 production by $CD4^+$ Tconv at day 15 following stimulation of WT and ΔUNG B16F10 tumor infiltrates with PMA/ionomycin. **(H)** Quantitation of IFN-$\gamma^+$TNF-$\alpha^+$, IFN-$\gamma^+$ (and TNF-$\alpha^{neg}$), or total IFN-$\gamma^+$ (TNF-$\alpha^{+/neg}$) tumor-infiltrating $CD8^+$ T cells, as percentages of the $CD45^+$ immune cell infiltrate. **(I)** Quantitation of IL-$17^+$ (Th17 cells), IFN-$\gamma^+$TNF-$\alpha^+$, and IFN-$\gamma^+$TNF-$\alpha^{neg}$ tumor-infiltrating $CD4^+$ Tconv, as percentages of the $CD45^+$ immune cell infiltrate. **(H-I)** Data from one experiment. n = 8 mice per group. **(D-I)** Mean ± SD bars shown. *$p<0.05$, **$p<0.01$ by two-tailed, unpaired t test. **(J-K)** Cell surface expression of MHC-I and PD-L1 were analyzed by flow cytometry on WT and ΔUNG B16F10 cells treated *in vitro* with IFN-γ (0.1, 0.3, or 1.0 ng/mL) for 18 h. Raw MFI were quantified and normalized to the mean of untreated WT cells within a given experiment. **(J)** Normalized MHC-I MFI for two independent repeat experiments, each with 3 biological replicates. **(K)** Normalized PD-L1 MFI combined from three independent experiments, each with 3 biological replicates. **(J-K)** Mean ± SD bars shown. ***$p<0.001$, ****$p<0.0001$ by two-way ANOVA with Tukey's multiple comparisons test.

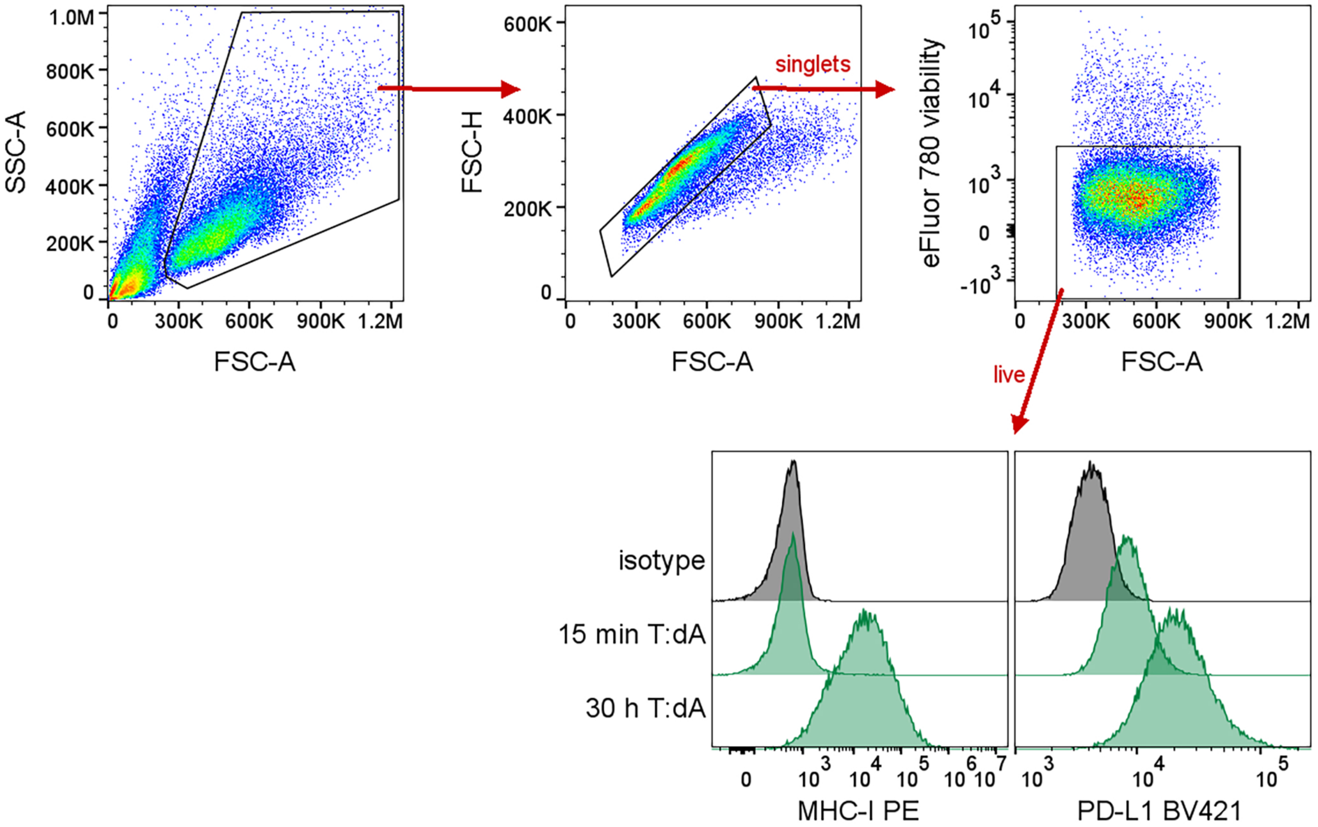


Pandya et al Figure S9

**Supplemental Figure S9. Gating strategy for MHC-I and PD-L1 cell surface expression *in vitro.***

Cells were gated based on FSC-A vs. SSC-A, followed by FSC-H vs. FSC-A, to exclude debris and doublets, respectively. Live cells (eFluor 780 viability dye negative) were analyzed for the relative surface expression of MHC-I and PD-L1, as measured by their median fluorescence intensity (MFI). Shown are representative histograms for MHC-I and PD-L1 expression following transfection with T:dA oligos for 15 min or 30 h, versus isotype control.

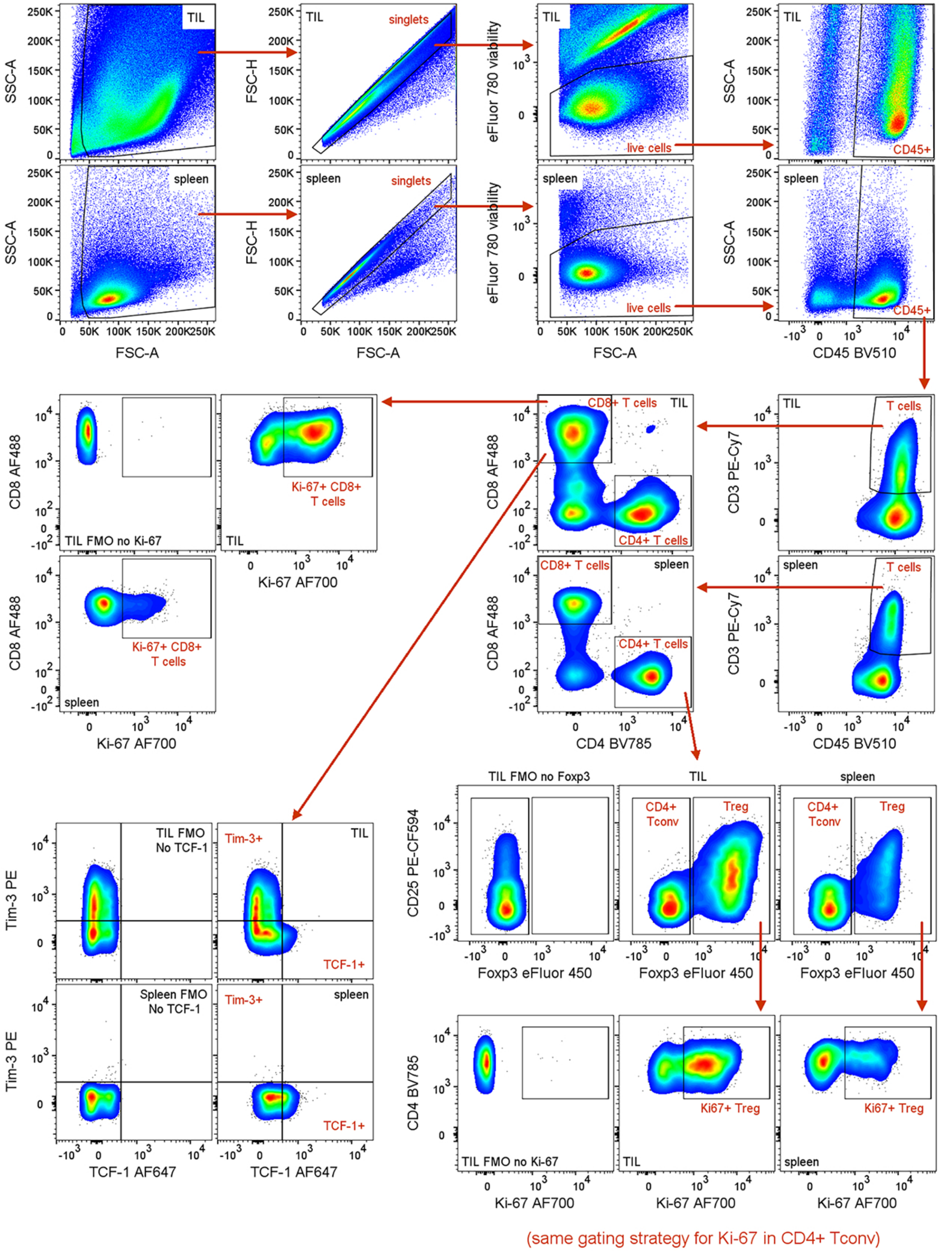


Pandya et al Figure S10

**Supplemental Figure S10. Gating strategy for profiling of tumor-infiltrating T cells.**

Any unstable portions of the run were excluded, and cells were gated based on FSC-A vs. SSC-A, followed by FSC-H vs. FSC-A, to exclude debris and doublets, respectively. Live (viability dye negative) cells were gated on CD45 expression, and $CD45^+$ immune cells were gated based expression CD3 to identify $CD3^+$ T cells, which were further subset into $CD8^+$ and $CD4^+$ T cells. $CD4^+$ T cells were further divided, based on expression of Foxp3 and CD25, into $Foxp3^{neg}$ conventional $CD4^+$ T cells ($CD4^+$ Tconv) and $Foxp3^+$ regulatory T cells (Treg). The marker CD25 was included to aid in resolution of the $CD4^+$ Tconv and Treg populations. TIL and spleen fluorescence minus one (FMO) controls (without Foxp3 eFluor 450) were included to determine gating for Foxp3. The $CD8^+$ T cell, $CD4^+$ Tconv, and Treg populations were all analyzed for expression of Ki-67. Gating for Ki-67 was determined using the spleen gating control sample in addition to FMO controls (without Ki-67 AF700). $CD8^+$ T cells were also examined for expression of Tim-3 and TCF-1, with gating determined using TIL and spleen FMO controls (without TCF-1 AF647) for the TCF-1 gating and the spleen gating control for the Tim-3 gate. Since no differences in these markers across groups were observed, quantitation was not included in the manuscript.

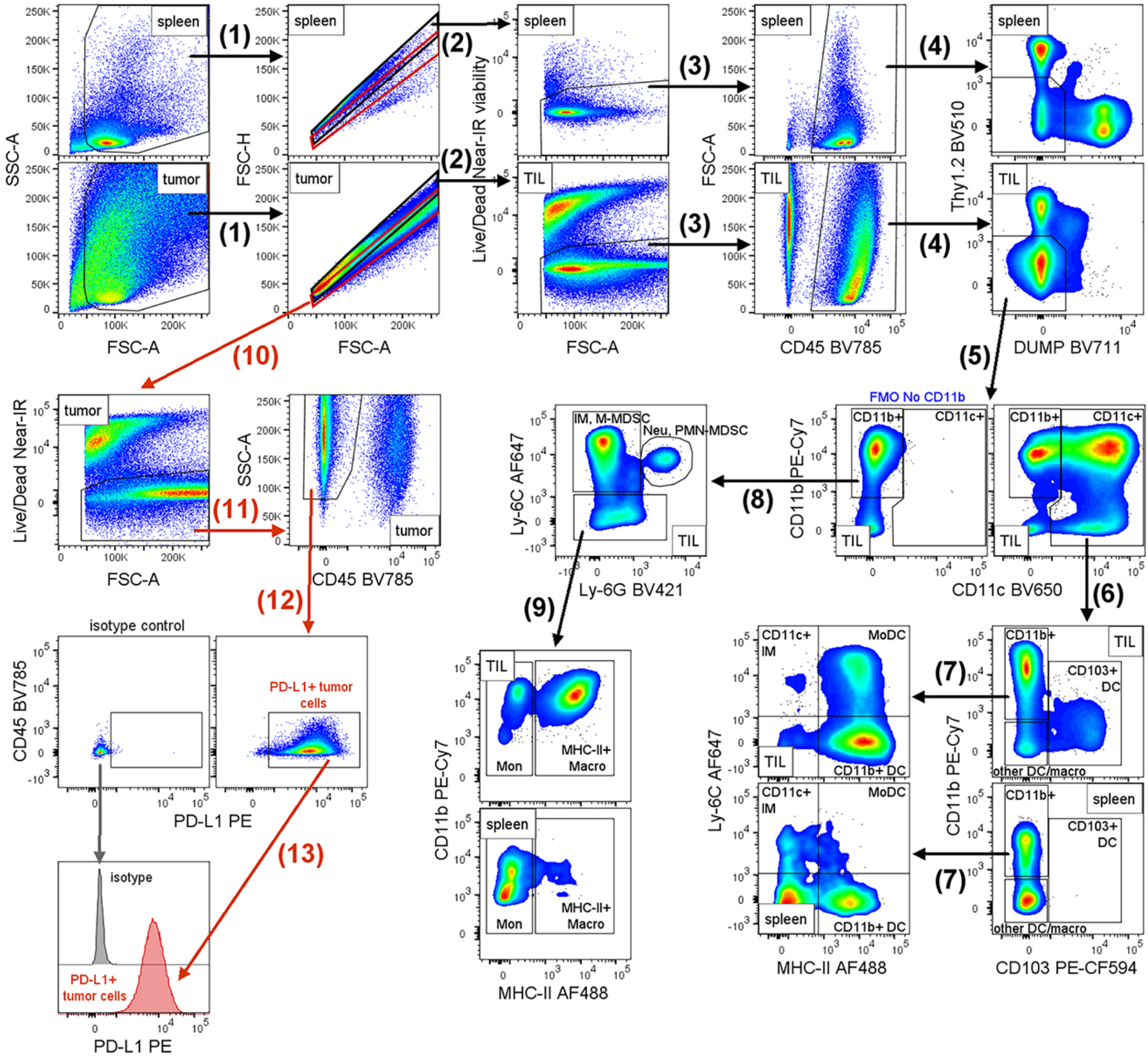


Pandya et al Figure S11

**Supplemental Figure S11. Gating strategy for profiling of tumor-infiltrating myeloid cells and tumor PD-L1 expression at day 14.**

Any unstable portions of the run were excluded, and cells were gated on FSC-A vs. SSC-A to remove debris, followed by **(1)** gating on FSC-H vs. FSC-A to remove doublets. Since immune cells and tumor cells have different FSC-H vs. FSC-A characteristics, two single cell gates were created. The **black** gate was drawn based on the spleen sample to identify the single cell immune population for myeloid profiling. The **red** gate was drawn to identify the single cell tumor cell population for analysis of tumor PD-L1 expression.

Immune cells within the **black single cell gate** were subset into myeloid populations. **(2)** Live cells were identified as Live/Dead Near-IR viability dye negative. **(3)** $CD45^+$ immune cells were gated. **(4)** $CD45^+$ cells negative for the lineage markers Thy1.2, CD19, and NK1.1 were gated to remove T cells, B cells and NK cells, respectively (CD19 and NK1.1 antibodies were combined in a BV711 DUMP channel). **(5)** Lineage negative cells were subset into total $CD11c^+$ ($CD11b^+$ or $CD11b^{neg}$) cells and $CD11b^+CD11c^{neg}$ cells. **(6)** Total $CD11c^+$ cells were subdivided based on expression of CD103 and CD11b. $CD11c^+CD103^+$ cells were defined as $CD103^+$ dendritic cells ($CD103^+$ DC), while $CD11c^+CD103^{neg}CD11b^{neg}$ were defined as other dendritic cells or macrophages (other DC/macro) without further characterization. **(7)** $CD11c^+CD11b^+CD103^{neg}$ cells were subdivided based on Ly-6C and MHC-II expression as follows: $CD11c^+CD11b^+$Ly-$6C^{neg}$MHC-$II^+$ cells were defined as $CD11b^+$ dendritic cells ($CD11b^+$ DC), $CD11c^+CD11b^+$Ly-$6C^+$MHC-$II^{neg}$ cells were defined as $CD11c^+$ inflammatory monocytes ($CD11c^+$ IM), and $CD11c^+CD11b^+$Ly-$6C^+$MHC-$II^+$ cells were defined as monocyte-derived dendritic cells (Mo-DC). **(8)** Lineage negative $CD11b^+CD11c^{neg}$ cells were subdivided based expression of Ly-6C and Ly-6C. $CD11b^+CD11c^{neg}$Ly-$6C^{hi}$Ly-$6G^{neg}$ were defined as inflammatory monocytes (IM) or monocytic (mononuclear) myeloid-derived suppressor cells (M-MDSC). $CD11b^+CD11c^{neg}$Ly-$6C^{int}$Ly-$6G^+$ were defined as neutrophils (Neu) or granulocytic (polymorphonucler) myeloid-derived suppressor cells (PMN-MDSC). **(9)** Cells negative for both Ly-6C and Ly-6G were subset based on MHC-II expression. $CD11b^+$MHC-$II^{neg}$ (and CD11c/Ly-6C/Ly-6G negative) cells were defined as monocytes ($CD11b^+$MHC-$II^{neg}$ Mon), while $CD11b^+$MHC-$II^+$ (and CD11c/Ly-6C/Ly-6G negative) were defined as MHC-$II^+$ macrophages ($CD11b^+$MHC-$II^+$ Macro).

Within the **red single cell gate**, **(10)** live cells were identified as Live/Dead Near-IR viability dye negative. **(11)** $CD45^{neg}$ tumor cells were gated. **(12)** $CD45^{neg}$ tumor cells were then gated based expression of PD-L1 (determined empirically using a fluorescence minus one (FMO) control without PD-L1 PE (not shown), and median fluorescence intensities (MFI) were determined. An isotype control tumor sample was included to verify minimal background signal. **(13)** The histogram depicts PD-L1 staining intensity for the PD-$L1^+$ population in a representative tumor versus background signal for isotype control staining on total $CD45^{neg}$ tumor population.

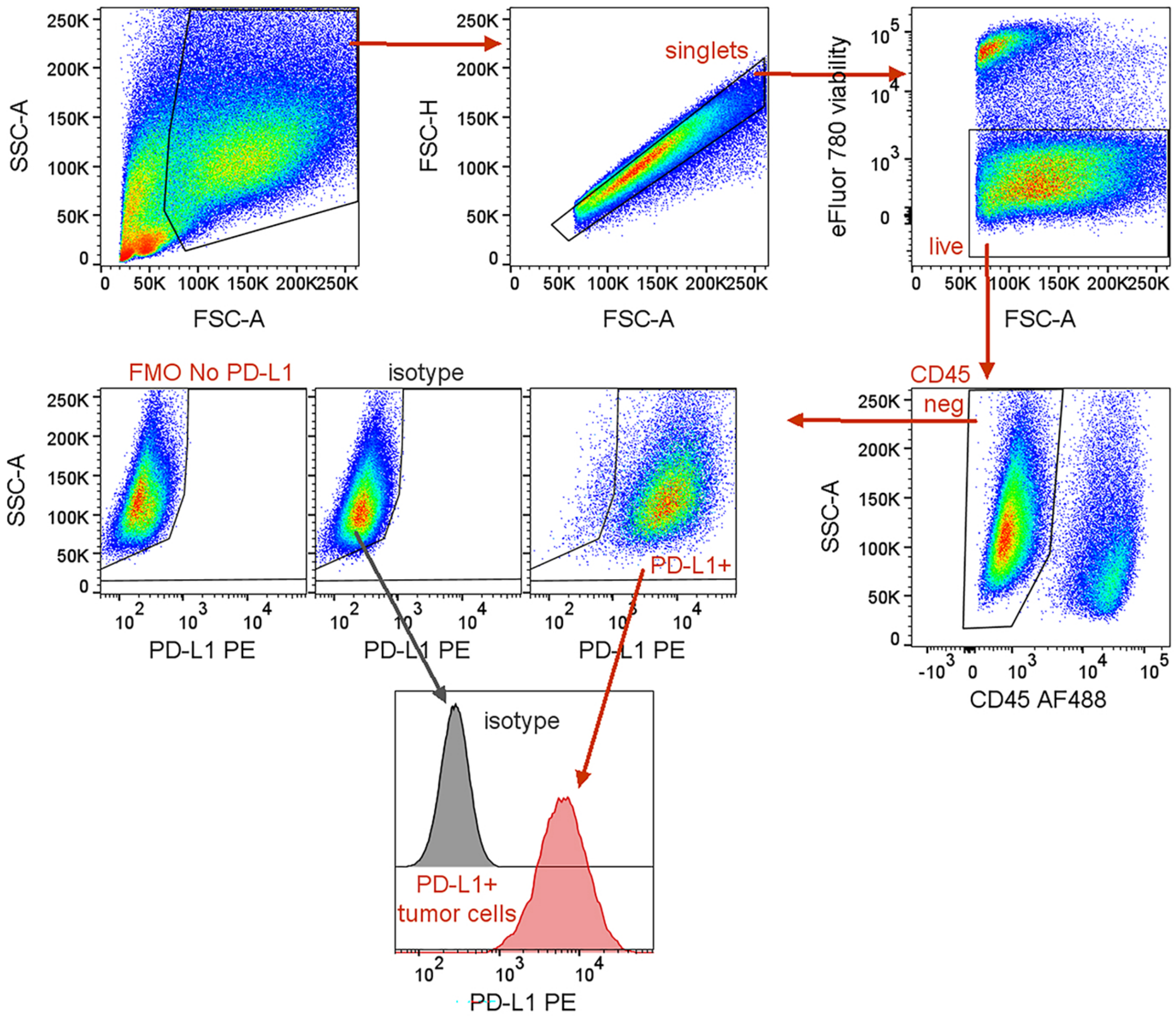


Pandya et al Figure S12

**Supplemental Figure S12. Gating strategy for profiling PD-L1 expression on tumors *in vivo* at day 15.** Any unstable portions of the run were excluded, and cells were gated based on FSC-A vs. SSC-A, followed by FSC-H vs. FSC-A, to exclude debris and doublets, respectively. Live (viability dye negative), $CD45^{neg}$ tumor cells were gated based expression of PD-L1 (determined empirically using a fluorescence minus one (FMO) control without PD-L1 PE, and median fluorescence intensities (MFI) were determined. The histogram depicts PD-L1 staining intensity for the $PD\text{-}L1^{+}$ population in a representative tumor versus background signal for isotype control staining on total $CD45^{neg}$ tumor population.

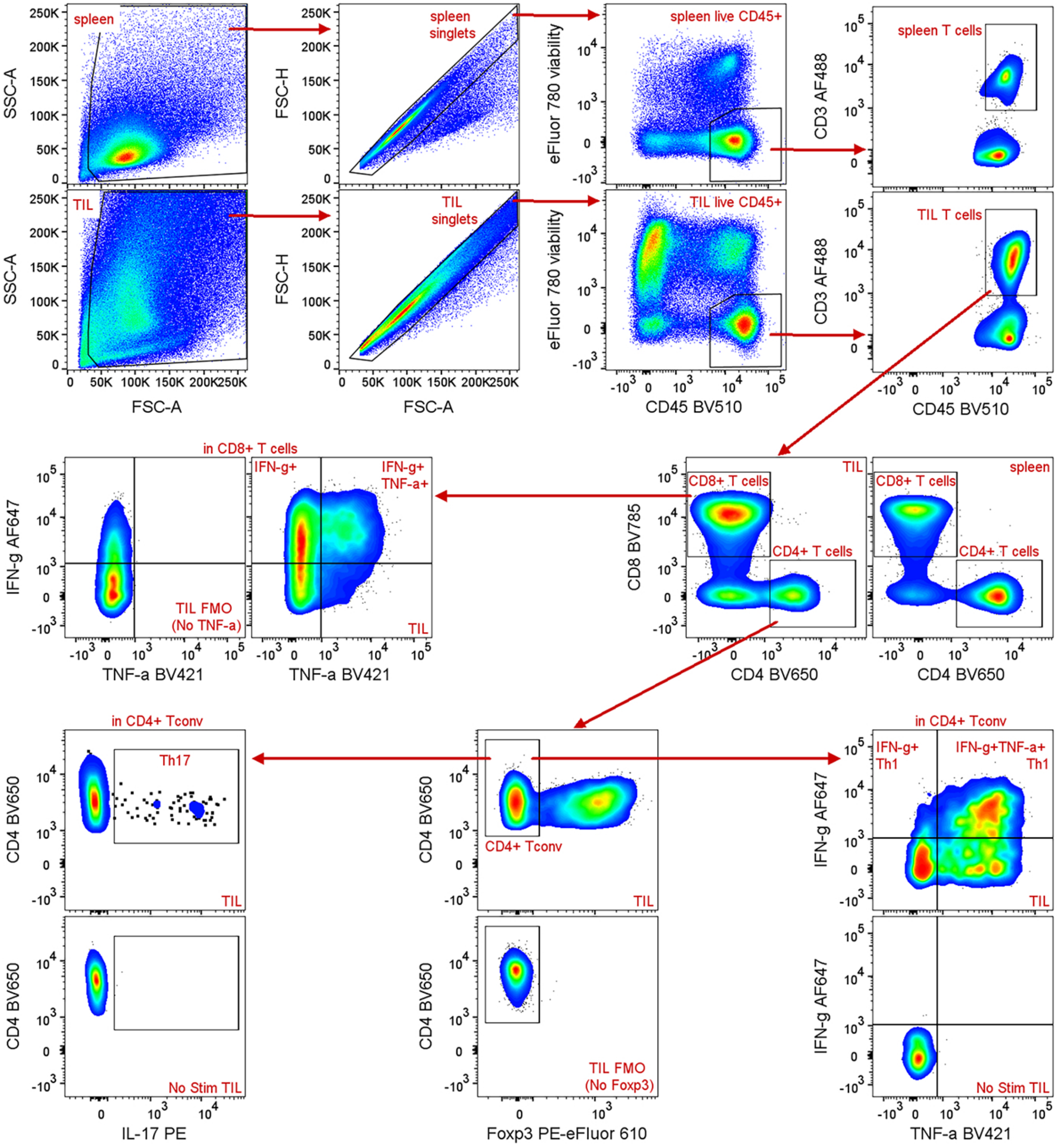


Pandya et al Figure S13

**Supplemental Figure S13. Gating strategy for profiling of tumor-infiltrating, cytokine-competent T cells.** Any unstable portions of the run were excluded, and cells were gated based on FSC-A vs. SSC-A, followed by FSC-H vs. FSC-A, to exclude debris and doublets, respectively. Live (viability dye negative), $CD45^+$ immune cells were gated, and total T cells ($CD3^+$) were divided into $CD8^+$ and $CD4^+$ T cell populations. $CD8^+$ T cells were examined for production of IFN-γ and TNF-α. $CD4^+$ T cells were examined for Foxp3 expression to exclude regulatory T cells from the conventional $CD4^+$ T cells ($CD4^+$ Tconv), which were examined for production of IFN-γ and TNF-α (to identify the Th1 subset) or IL-17 (to identify the Th17 subset). TIL and spleen unstimulated (No Stim) controls and No TNF-α fluorescence minus one (FMO) controls were used to determine gating for IFN-γ and TNF-α in both $CD8^+$ T cells and $CD4^+$ Tconv. No Foxp3 FMO controls were used to determine gating for $CD4^+$ Tconv (ie. $Foxp3^{neg}$). IL-17 gating was determined using the No Stim controls.

### Key Resources Table

**Western Blot**

| Reagent | Source | Identifier |
|---|---|---|
| TBK1 (1:1000 Dilution) | Cell Signaling Technology | 3504 |
| Phospho-TBK1 (ser172) (1:1000 Dilution) | Cell Signaling Technology | 5483 |
| Phospho-STAT (Tyr701) (1:1000 Dilution) | Cell Signaling Technology | 9167 |
| GAPDH (1:50000 Dilution) | Abcam | Ab9485 |

**Flow Cytometry**

| Reagent | Source | Identifier |
|---|---|---|
| AF647 anti-mouse TCF-7/TCF-1, clone S33-966 (1:200 Dilution) | BD Biosciences | 566693 |
| BV421 anti-mouse CD274 (PD-L1), clone 10F.9G2 (1:100 Dilution) | BD Biosciences | 568309 |
| BV421 rat IgG2b, κ isotype control, clone R35-38 (1:100 Dilution) | BD Biosciences | 562603 |
| BV650 anti-mouse CD4, clone GK1.5 (1:500 Dilution) | BD Biosciences | 563747 |
| PE-CF594 anti-mouse CD25, clone PC61 (1:250 Dilution) | BD Biosciences | 562695 |
| PE-CF594 anti-mouse CD103, clone M290 (1:250 Dilution) | BD Biosciences | 565849 |
| AF488 anti-mouse CD3, clone 17A2 (1:250 Dilution) | BioLegend | 100212 |
| AF488 anti-mouse CD8α, clone 53-6.7 (1:250 Dilution) | BioLegend | 100723 |
| AF488 anti-mouse CD45, clone 30-F11 (1:250 Dilution) | BioLegend | 103122 |
| AF488 anti-mouse I-A/I-E (MHC-II), clone M5/114.15.2 (1:500 Dilution) | BioLegend | 107615 |
| AF647 anti-mouse IFN-γ (clone XMG1.2) (1:500 Dilution) | BioLegend | 505816 |
| AF647 anti-mouse Ly-6C, clone HK1.4 (1:500 Dilution) | BioLegend | 128010 |
| AF700 anti-mouse Ki67, clone 16A8 (1:200 Dilution) | BioLegend | 652420 |
| BV421 anti-mouse Ly-6G, clone 1A8 (1:250 Dilution) | BioLegend | 127628 |
| BV421 anti-mouse TNF-α, clone MP6-XT22 (1:250 Dilution) | BioLegend | 506328 |
| BV510 anti-mouse CD45, clone 30-F11 (1:250 Dilution) | BioLegend | 103138 |
| BV510 anti-mouse Thy1.2 (CD90.2), clone 30-H12 (1:250 Dilution) | BioLegend | 105335 |
| BV650 anti-mouse CD11c, clone N418 (1:100 Dilution) | BioLegend | 117339 |

| | | |
|---|---|---|
| BV711 anti-mouse CD19, clone 6D5<br>(1:250 Dilution) | BioLegend | 115555 |
| BV711 anti-mouse NK-1.1, clone PK136<br>(1:250 Dilution) | BioLegend | 108745 |
| BV785 anti-mouse CD4, clone GK1.5<br>(1:500 Dilution) | BioLegend | 100453 |
| BV785 anti-mouse CD8α, clone 53-6.7<br>(1:250 Dilution) | BioLegend | 100750 |
| BV785 anti-mouse CD45, clone 30-F11<br>(1:250 Dilution) | BioLegend | 103149 |
| PE anti-mouse IL-17A, clone TC11-18H10.1<br>(1:200) | BioLegend | 506903 |
| PE-Cy7 anti-mouse/human CD11b, clone M1/70<br>(1:250 Dilution) | BioLegend | 101215 |
| PE anti-mouse CD274 (PD-L1), clone 10F.9G2<br>(1:100 Dilution *in vitro*, 1:250 Dilution *in vivo*) | BioLegend | 124307 |
| PE anti-mouse CD366 (Tim-3), clone RMT3-23<br>(1:250 Dilution) | BioLegend | 119703 |
| PE anti-mouse H-2Ld/H-2Db, clone 28-14-8<br>(1:100 Dilution) | BioLegend | 114507 |
| PE mouse IgG2a, κ isotype control, clone MOPC-173<br>(1:100 Dilution) | BioLegend | 400211 |
| PE rat IgG2b, κ isotype control, RTK4530<br>(1:100 Dilution *in vitro*, 1:250 Dilution *in vivo*) | BioLegend | 400607 |
| PE-Cy7 anti-mouse CD3, clone 17A2<br>(1:250 Dilution) | BioLegend | 100219 |
| eFluor 450 anti-mouse/rat Foxp3, clone FJK-16s<br>(1:200 Dilution) | Invitrogen | 48-5773-82 |
| PE-eFluor 610 anti-mouse/rat Foxp3, clone FJK-16s<br>(1:200 Dilution) | Invitrogen | 61-5773-80 |
| BD Horizon™ Brilliant Stain Buffer Plus<br>(1:5 Dilution) | BD Biosciences | 566385 |
| TruStain FcX™ PLUS (anti-mouse CD16/32) Antibody<br>(1:100 Dilution) | BioLegend | 156604 |
| True-Stain Monocyte Blocker™<br>(1:20 Dilution) | BioLegend | 426103 |
| FluoroFix buffer | BioLegend | 422101 |
| eBioscience Protein Transport Inhibitor Cocktail<br>(500X) | Invitrogen | 00-4980-93 |
| eBioscience Cell Stimulation Cocktail (plus protein transport inhibitors) (500X) | Invitrogen | 00-4975-03 |
| eBioscience fixable viability dye eFluor 780<br>(1:2000-1:3000 Dilution) | Invitrogen | 65-0865-18 |
| eBioscience normal mouse serum<br>(used at 5% in 1x PBS) | Invitrogen | 24-5544-94 |
| eBioscience flow cytometry staining (FCS) buffer | Invitrogen | 00-4222-26 |
| eBioscience Permeabilization Buffer (10X) | Invitrogen | 00-8333-56 |

| | | |
|---|---|---|
| eBioscience Fixation/Permeabilization Concentrate | Invitrogen | 00-5123-43 |
| eBioscience Fixation/Permeabilization Diluent | Invitrogen | 00-5223-56 |
| LIVE/DEAD™ Fixable Near-IR dead cell stain (1:1000 Dilution) | Invitrogen | L10119 |
| OneComp eBeads compensation beads | Invitrogen | 01-1111-42 |
| Collagenase Type IV, Filtered | Worthington | LS004209 |
| Deoxyribonuclease I | Worthington | LS002139 |
| Trypsin Inhibitor, Soybean (Animal-Free) | Worthington | LS003587 |

**Transfection**

| Reagent | Source | Identifier |
|---|---|---|
| Cy3 siRNA transfection control | Invitrogen | AM4621 |
| Poly(dA:dT)/Lyovec™ | Invitrogen | tlrl-patc |
| LyovecTM | Invitrogen | Lyec-1 |
| DharmaFECT1 transfection reagent | Horizon discovery | |
| TransIT-X2 Dynamic Delivery System | mirusbio | MIR 6004 |
| Lenti-XTM Concentrator | Takarabio | 631231 |
| LipofectamineTM CRISPRMAXTM cas9 transfection reagent | Thermofisher | CMAX00003 |
| Opti-MEMTM I reduced serum | Thermofisher | 31985062 |
| Gene knockout kit v2 mouse-Tmem173 (sting)-3.0nm (Crispr guide) | Synthego | |
| SpCas9, 1000pmol | Synthego | |

**Recombinant DNA**

| Plasmid | Source | Identifier |
|---|---|---|
| pMDLg/pRRE | Addgene | 12251 |
| pRSV-Rev | Addgene | 12253 |
| pMD2.G | Addgene | 12259 |
| pLENTI-CRISPR-V2-control gRNA_1 | This study | Sobol Lab 2014 |
| pLENTI-CRISPR-V2-control gRNA_2 | This study | Sobol Lab 2015 |
| pLV-hCas9-T2A-Puro-U6-mUng-gRNA9 | This study | Sobolo Lab 2416 |

**Chemicals/Inhibitors**

| Reagent | Source | Identifier |
|---|---|---|
| AZD6738 (ATRi) | AstraZeneca | |
| AZD0156 (ATMi) | AstraZeneca | |
| LNT 1 (Fen1i) | AstraZeneca | 6510 |
| 2’3’-cGAMP | Medchem Express | HY-100564 |
| ML-60218 (RNA polymerase-III inhibitor) | Medchem Express | HY-122122 |
| Recombinant Murine IFN-γ (Animal-Free) | Peprotech | AF-315-05 |
| inVivoPlus anti-mouse PD-L1 (B7-H1) | BioXCell | BP0101 |
| inVivoPure pH 6.5 Dilution Buffer | BioXCell | IP0065 |

| | | |
|---|---|---|
| Puromycin | InvivoGen | ant-pr-1 |
| RPMI-1640 | Lonza | 12-702F |
| DMEM | Lonza | 12-604F |
| Phosphate buffered saline (PBS) | Fisher | BP2944-100 |
| Ethanol | Decon Labs, Inc. | 2716 |
| Proteinase K | Invitrogen | AM2546 |
| RNase A | Thermo Scientific | EN0531 |
| RNase H | New England BioLabs | M0297 |
| RNase H Buffer | New England BioLabs | B0297 |
| RNase/DNase free water | MP Biomedicals | 8217739 |
| Hind III | Thermo Scientific | FD0504 |
| EcoRI | Thermo Scientific | FD0274 |
| Bam HI | Thermo Scientific | FD0054 |
| FastDigest Buffer (10X) | Thermo Scientific | B64 |
| DNA Degradase Plus | Zymo Research | E2021 |
| 10X DNA Degradase Reaction Buffer | Zymo Research | E2016-2 |
| Trypan Blue Solution (0.4%) | Sigma | T8154 |
| Fetal Bovine Serum (FBS) | GEMINI Bioproducts | 900-108 |
| 0.05% Trypsin-EDTA (1x) | Gibco | 25300-054 |
| Pen/Strep | Gibco | 15140-122 |
| MEM NEAA | Gibco | 11140-050 |
| Sodium pyruvate | Gibco | 11360-070 |
| HEPES | Gibco | 15630-080 |
| HEPES | Corning | 25060CI |
| β-mercaptoethanol | Gibco | 21985-023 |
| 0.5 M EDTA pH 8.0 | Boston Bio Products, Inc. | BM-150 |
| Optima LC-MS grade water | ThermoFisher | W6-4 |
| Optima LC-MS grade acetonitrile | ThermoFisher | A955-4 |
| Optima LC-MS grade methanol | ThermoFisher | A456-4 |
| 1,1,1,3,3,3-hexafluoro 2-propanol (HFIP) | Sigma-Aldrich | 105228 |
| Diisopropylethylamine (DIPEA) | Sigma-Aldrich | D125806 |
| Dimethyl sulfoxide (DMSO) | Fisher | BP231-1 |
| Tris-HCl pH8.0 | Invitrogen | 15568-025 |
| Tris-HCl pH 7.5 | Invitrogen | 15567-027 |
| NaOH | Sigma-Aldrich | S8045-500G |
| NaCl | Fisher | S271-10 |
| Mouse serum | MP Biomedicals | 152282 |
| Whatman Hybond N+ Blotting Membrane | Millipore Sigma | Z761079 |
| Triton X100 | Fisher | BP151-500 |
| Digitonin | Cayman Chemical | 14952 |
| 10 % sodium dodecyl sulfate (SDS) | Fisher | BP2436-1 |
| Leammli buffer | BioRad | 1610737 |
| 16% (w/v) Paraformaldehyde (PFA) | Fisher | AA433689M |
| Tween-20 | Fisher | BP337-500 |
| Glycerol | EMD | GX0185-5 |
| NP40 | Fluka BioChemika | 56741 |

| | | |
|---|---|---|
| cOmplete Mini, EDTA-free (protease inhibitor) | Roche | 11836170001 |
| Trizol | Ambion by Life Technologies | 15596026 |
| Bovine Serum Albumin (BSA) DNase- and Protease-free Powder | Fisher BioReagents | BP9706100 |

**Immunofluorescence**

| Reagent | Source | Identifier |
|---|---|---|
| Anti-Rabbit cGAS (D3O8O) (1:200) | Cell Signaling | 31659 |
| Goat anti-Rabbit IgG (H+L) Alexa Fluor™ 488 (2 µg/ml) | Invitrogen | A32731 |

**In Vitro BER**

| Reagent | Source | Identifier |
|---|---|---|
| Uracil DNA Glycosylase (5U) | New England BioLabs | M0280S |
| Endonuclease IV (15U) | New England BioLabs | M0304S |
| Bst full-length polymerase (4U) | New England BioLabs | M0328S |
| Taq Ligase (68U) | New England BioLabs | M0208S |
| NAD+ (500 µM) | Thermo Scientific Chemicals | J62337.06 |
| dATP (200 nM) | Invitrogen | R0141 |
| dCTP (200 nM) | Invitrogen | R0151 |
| dGTP (200 nM) | Invitrogen | R0161 |
| Cy3-dUTP (200 nM) | Jena Biosciences | NU-803-CY3-L |
| Paraformaldehyde (2%) | Thermo Scientific Chemicals | 30525-89-4 |
| Triton-X 100 (0.5%) | Fisher BioReagents™ | BP151-100 |
| Ultrapure Sucrose (300 mM) | Bethesda Research Laboratories | 5503UA |
| NaCl (100 mM) | Sigma-Aldrich | S7653-250G |
| MgCl2 (3 mM) | Invitrogen | AM9530G |
| PIPES (10 mM) | Thermo Scientific Chemicals | A16090.22 |
| Fluoroshield with DAPI | Sigma Aldrich | F6057 |

**Commercial Kits**

| Kit | Source | Identifier |
|---|---|---|
| Quick-DNA Miniprep Plus kit | Zymo research | D4069 |
| Direct-zol RNA isolation kit | Zymo research | R2052 |
| Lunascript RT master mix | NEB | M3010 |
| Luna Universal qPCR master mix | NEB | M3003 |
| PowerUp™ SYBR™ Green Master Mix | Applied Biosystems | A25742 |
| CellTiter-Glo® 2.0 Cell viability assay | Promega | G9242 |
| NucBusterTM protein extraction kit | Millipore | 71183 |
| PureLink Genomic DNA Mini Kit | Invitrogen | K182002 |
| GeneJET PCR Purification Kit | Thermo Scientific | K0702 |

**Experimental Models (Cell lines/Mouse)**

| | | |
|---|---|---|
| 293-FT (derived from human embryonal kidney cells transformed with the SV40 large T antigen) | Thermofisher Scientific | R70007 |
| Mouse: B16F10 melanoma cell line | ATCC | CRL-6475 |
| B16F10 Cas9 | This study | |
| B16F10/ ΔUNG | This study | |
| B16F10/ ΔUNGΔPP | This study | |
| Mouse: YUMM1.7 melanoma cell line | Aird Lab | |
| YUMM1.7/*ung-/-* polyclonal | This study | |
| Mouse: MEF 92Tag (WT) | Sobol Lab1 | Clone# 5689 |
| MEF 207Tag (UNG KO) | Sobol Lab2 | Clone#7406 |
| MEF 88Tag (Polβ-KO) | Sobol Lab1 | Clone#5689 |
| Mouse: C57BL/6J | Jackson | JAX:000664 |
| Mouse: athymic nude mice (NU/J) | Jackson | JAX:002019 |

**qPCR Primer**

IFNβ Forward: 5'-AAGAGTTACACTGCCTTTGCCATC-3’

IFNβ Reverse: 5'-CACTGTCTGCTGGTGGAGTTCATC-3’

GAPDH Forward: 5´-CTCTGGAAAGCTGTGGCGTGATG-3’

GAPDH Reverse: 5´-ATGCCAGTGAGCTTCCCGTTCAG-3’

DDX58 Forward: 5’-GAGACCGAGCGAGAGCTTAC-3'

DDX58 Reverse: 5’-CTGTTGCCCGGTTTGTTCTG-3'

IFIH1 Forward: 5’-CTATCCCGTGTGGCTGGTAG-3'

IFIH1 Reverse: 5’-CTCCAAGATTCCTTCCCAAA-3'

Sting Forward: 5’-GGAACACCGGTCTAGGAAGC-3'

Sting Reverse: 5’-TGGATCCTTTGCCACCCAAA-3'

Trim 30a Forward: 5’-CCTGATCGTCCCAGTCGTGT-3’

Trim 30a Reverse: 5’-AGTGCCAGCTTTCCGATCC-3'

PARP12 Forward: 5’-CTGGAGCAGTTGGAAAGGTTGGG-3'

PARP12 Reverse: 5’-GCGGGAGAAGGAGACACTTTGC-3'

RNF213 Forward: 5’-TTTGTACCGTTCCCCCAAT-3'

RNF213 Reverse: 5’-GTTCACTGCCTCCAATTGCT-3'

Gusb Forward: 5’-GTAAAGAATACGTGGTCGGAGAG-3’

Gusb Reverse: 5’-AGTTTTGGGCTGTCTCTGG-3’

Tubb5 Forward: 5’-GTTCTGGGAGGTGATAAGCG-3’

Tubb5 Reverse: 5’-GATAGCTCGAGGGACATACTTG-3’

**Primers used for Sanger Sequence**

Pair 1 Forward: GTTAAATCAGGTTGGCGGGCTG

Pair 1 Reverse: GGCCATCCCTTCTAGTCTGTGC

Pair 2 Forward: GAAGTCAGGAAAGACGATCGAGGC

Pair 2 Reverse: CGTTGCCTACACTCAATAGACTGAG

**Oligonucleotides/gRNA for Molecular Beacon Assay**

| Sequence | Company | Lesion |
|---|---|---|
| 6-Fam-CCA CTA TTG AAT TGA CAC GCC ATG TCG ATC AAT TCA ATA GTG G-Dabcyl | IDT | FD-Con2[7] |
| 6-Fam-CCA CTX TTG AAT TGA CAC GCC ATG TCG ATC AAT TCA ATA GTG G-Dabcyl | IDT | FD-THF2,where X indicates the location of lesion[7] |
| 6-Fam-CCA CTX TTG AAT TGA CAC GCC ATG TCG ATC AAT TCA AAA GTG G-Dabcyl | IDT | FD-dU/A, where X indicates the location of the lesion[7] |
| 6-Fam-CCA CTX GTG AAT TGA CAG CCC ATG TGC ATC AAT TCA CGA GTG G-Dabcyl | IDT | FDB-dU/G, where X indicates the location of the lesion[7] |
| 6-Fam-CCA CCG TTG AAT TGA CAG CCC ATG TGC ATC AAT TCA AXG GTG G-Dabcyl | IDT | FDB-G/dU, where X indicates the location of the lesion[7] |

| **dsDNA fragments for Transfection** | | |
|---|---|---|
| Thymidine Sting Agonist - Sense | CAAAGTGAGTTCCAGGACAGTCAGGGCTACACAGAGAAACC<br>CTGTCTCGGAAAAAAAAAAAAAAAAAAAAAAAAAAAACTCA<br>AAAAAGAAAAAAAAAAAAAGATTGGTCTGTGGGGTTCCTTT<br>TTTTGTTTTTTCGAGACAGTGTTTCTCTGTGTAGCCCTGGCTG<br>TCCTGGAACTCACTCTGTAGACCAGGCTGGCCTC | IDT |
| Thymidine Sting Agonist - Antisense | GAGGCCAGCCTGGTCTACAGAGTGAGTTCCAGGACAGCCAG<br>GGCTACACAGAGAAACACTGTCTCGAAAAAACAAAAAAAGG<br>AACCCCACAGACCAATCTTTTTTTTTTTTTCTTTTTTGAGTTTTT<br>TTTTTTTTTTTTTTTTTTTTTTTCCGAGACAGGGTTTCTCTGTGT<br>AGCCCTGACTGTCCTGGAACTCACTTTG | IDT |
| dU Sting Antagonist - Sense | CAAAGTGAGTTCCAGGACAGTCAGGGCTACACAGAGAAACC<br>CTGTCTCGGAAAAAAAAAAAAAAAAAAAAAAAAAAAACTCA<br>AAAAAGAAAAAAAAAAAAAGATTGGTCTGTGGGGTTCCTTT<br>TTTTGTTTTTTCGAGACAGTGTTTCTCTGTGTAGCCCTGGCTG<br>TCCTGGAACTCACTCTGTAGACCAGG | IDT |
| dU Sting Antagonist - Antisense | GAGGCCAGCCTGGTCTACAGAGTGAGTTCCAGGACAGCCAG<br>GGCTACACAGAGAAACACTGTCTCGAAAAAACAAAAAAAGG<br>AACCCCACAGACCAATCUUUUUTTTTTTTTTCTTTUUTGAGUU<br>UUUTTTTTTTTTTTTTTTTTTTUUUUUCCGAGACAGGGUUUCT<br>CTGTGTAGCCCTGACTGTCCTGGAACTCACTTTG | IDT |
| Thymidine RNA polymerase III agonist - Sense | TATATATGTATATATATGTATATATATGTATATACATATATAT<br>GTATATATATGTATATATATGTGTATATATACATATATATGTA<br>TATATATGTATATATATATGTATATATATGTATATATATACAT<br>ATATATGTATATATATGTATATATATACATATATATGTATATA<br>TATGTATATATATATGTATGTATATATA | IDT |

| | | |
|---|---|---|
| Thymidine RNA polymerase III agonist - Antisense | TATATATACATACATATATATATACATATATATACATATATATGTATATATATACATATATATACATATATATGTATATATATACATATATATACATATATATATACATATATATACATATATATGTATATATACACATATATATACATATATATACATATATATGTATATACATATATATACATATATATACATATATA | IDT |
| dU RNA Polymerase Antagonist - Sense | TATATATGTATATATATGTATATATATGTATATACATATATATGTATATATATGTATAUAUAUGTGTAUAUAUACATATATATGUAUAUAUATGTATATATATATGTAUAUAUATGTAUAUAUATACATATATATGUAUAUAUATGTATATATATACATATATATGTATATATATGTATATATATATGTATGTATATATA | IDT |
| dU RNA Polymerase Antagonist - AntiSense | TATATATACATACATATATATATACATATATATACATATATATGTATATATATACAUAUAUAUACATATATATGTAUAUAUATACATAUAUAUACATATATATATACAUAUAUAUACATATATATGUAUAUATACACAUAUAUATACATATATATACATATATATGTATATACATATATATACATATATATACATATATA | IDT |

**Software and algorithms**

| Resource | Source | Link |
|---|---|---|
| GraphPad Prism 10.1.1 | GraphPad | https://www.graphpad.com/scientific-software/prism/ |
| Quant Studio | Thermofisher | |
| Gene set enrichment analysis 4.3.2 | www.gsea-msigdb.org | |
| FlowJo v10 | BD Biosciences | https://www.flowjo.com/solutions/flowjo |
| ImageJ | Schneider et al., 2012 | https://imagej.nih.gov/ij/index.html |
| Adobe Illustrator 2023 | Adobe | https://www.adobe.com/products/illustrator.html |
| Adobe Photoshop Elements 15 | Adobe | https://www.adobe.com/products/photoshop.html |
| BioRender | BioRender | BioRender.com |
| NIS-Elements 5.3 | Nikon | https://www.microscope.healthcare.nikon.com/products/software/nis-elements |
| LCMS | | |

### Lead contact

### METHODS DETAILS

#### Cell Lines

Mouse embryonic fibroblasts (MEF) (WT 92Tag, *polb-/-* knockout 88Tag, *ung-/-* knockout 207Tag) were isolated from 14.5-day-old embryos and transformed by SV40 T-antigen expression (1,2). Each were isolated from 14.5-day-old embryos and transformed by SV40 T-antigen expression, as previously described. No drug resistance gene was inserted. MEF were grown in D-MEM supplemented with 10% heat-inactivated fetal bovine serum (FBS), 1% Pen/Strep and 5% GlutaMAX. 293-FT cells, obtained Thermo Fischer Scientific, were derived from human embryonal kidney cells and are transformed with the SV40 large T antigen. B16F10 cells used in this study include ΔUNG, ΔUNGΔPP, and ΔSTING. B16F10 cells were grown in D-MEM, supplemented with 10% heat-inactivated FBS, 1% Pen/Strep, and 1% GlutaMAX. YUMM1.7 cells used in this study include the YUMM1.7, YUMM1.7 Cas9 control and YUMM1.7 *ung-/-* knockout polyclonal. YUMM1.7 were grown in DMEM/F12 supplemented with 10% heat-inactivated FBS, 0.5% Pen/Strep and 5.6mL MEM Non-Essential Amino Acids. The media for the B16F10 and YUMM1.7 modified cells was supplemented with 1.5 µg/mL of puromycin. We routinely validate cell lines by Genetica Cell Line Testing. Mycoplasma contamination was monitored quarterly using Lonza MycoAlert (Lonza #LT07-318). Cells under passage number 15 were used for all experiments. All cells were grown in a humidified cell culture incubator set to 37°C, 5% $CO_2$.

#### Cell line treatment

Cell lines were treated with AZD6738 (ATRi) or AZD0156 (ATMi) in complete media. MEF were treated with 2.5um ATRi in the presence or absence of 5uM FEN1i for 24h.

For cGAMP stimulation, cells were treated with 10ug/ml cGAMP in digitonin permeabilization solution (50mM HEPES, pH 7.0, 100 mM KCL, 3mM $MgCl_2$, 0.1mM DTT, 85 mM sucrose, 0.2% BSA, 1 mM ATP, 0.1 mM GTP, and 10 ug/ml digitonin) at 37°C for 10min. Cells were then incubated in fresh media for 4 h before the cells were harvested for RNA isolation.

#### Commercial or synthesized dsDNA fragment transfection

Poly(dA:dT)/Lyovec™ was purchased from Invivogen and dissolved into 50 µg/ml using nuclease-free water. Cells were exposed to 0.2 ug/ml for 24h.

dsDNA fragments were synthesized by IDT. They were complexed with lyovec™ (Invivogen) (3,4). Cells were exposed to the complex at a final concentration of 0.1 ug/ml.

#### Lentivirus production and cell transduction

Virus production: 293-FT cells were used to prepare lentivirus expressing Cas9 and Cas9/sgRNA specific to mouse Ung. Cells ($1x10^6$/dish) were seeded in a 60mm dish for overnight incubation. Packaging vectors for the third-generation system pMDLg/pRRE (Addgene, Cat# 12251), pRSV-

Rev (Addgene, Cat# 12253), and pMD2.G (Addgene, Cat# 12259) and the shuttle vectors/transfer vectors (see Table S1) were co-transfected into 293-FT cells using the TransIT-X2 Dynamic Delivery System (Cat# MIR 6003). Supernatant containing the lentivirus was collected after 48 hours, followed by filtration using 0.45μM filters to remove cell debris and isolate the viral particles, as described previously(5-7). The lentivirus particles were then further concentrated using the Lenti-X Concentrator (Takara Bio, Cat# 631231), as per the manufacturer's instructions.

Transduction: Target cells ($2\times10^5$/well) were seeded into a six-well plate and cultured for 24 hours. The media was then replaced with 1 mL of complete fresh media followed by dropwise addition of the lentiviral particle solution (1 mL). After overnight incubation in the incubator, lentivirus-containing media was replaced with complete media with puromycin (1.5ug/ml) and kept for selection for 5 days. Fresh complete media was added on every alternative day.

Generation of UNG knockout cells: After puromycin selection, the polyclonal population was seeded into 96 well plates as single cells by serial dilution method. Once cell clones reached optimum confluency, they were harvested. Genomic DNA was isolated and amplified using primer pair 1 (see table UNG primer). Next, amplified DNA was purified using a PCR purification kit (ThemoFisher). Purified DNA with primer pair 2 (See table UNG primer) was submitted to Azenta Life Science (New Jersey, USA) for sanger sequence analysis to validate the knockout clone.

**Cell lysate preparation for the DNA repair molecular beacon assay**

Harvested cells were centrifuged at 500g, 4°C, for 5 minutes. The supernatant was discarded and the nuclear lysate from the cell pellets was then extracted using the NucBuster Protein Extraction Kit. All NucBuster steps were performed on ice. The protein content of the resulting nuclear protein extract was measured on a Nanodrop 2000 Spectrophotometer. All nuclear lysates were diluted to 2μg/uL in preparation for the DNA Repair Molecular Beacon (DRMB) assay. BER buffer was used to dilute the nuclear lysates.

**DNA repair molecular beacon assay**

The DRMB assay was performed as described previously (7). Details are as follows:

Buffer Preparation: The DRMB assay is run in a standard Base Excision Repair (BER) Reaction buffer that is comprised of 25mM HEPES-KOH, 150mM KCl, 0.5mM EDTA, 2% Glycerol, and 0.5mM DTT in autoclaved water. BER buffer was filtered using a 0.45μm filter.

Beacon and Annealing: DNA repair molecular beacons were annealed prior to use in the beacon assay to ensure the integrity of a hairpin structure. Five DNA repair molecular beacons were used: CON2, THF2, dU/dA, dU/dG, and dG/dU (see **Key Resources**). All DNA repair molecular beacons contain a 6-Fam fluorophore on the 5' end and a Dabcyl non-fluorescent quencher on the 3' end. CON2 contains no such modifications and is used as a negative control. THF2 contains tetrahydrofuran; mimicking an abasic sight and targeting APE1. The THF2 beacon serves as a positive control. The dU/A probe contains deoxyuridine opposite adenine. The dU/G probe contains deoxyuridine opposite guanine. The G/dU also contains a deoxyuridine opposite guanine, but in the reverse order. All beacons were diluted to 200nM using BER Reaction buffer into light-blocking microcentrifuge tubes. After dilution, beacons were placed in boiling water to denature the beacons. After 3 minutes, the heat source was removed from the boiling water and the beacons were left overnight to aneal into hairpin structures.

Assay: All samples were load into a MicroAmp Fast Optical 96-Well Reaction Plate on ice. Two sets of samples were loaded into the plate: experimental and control. For each experimental sample: 15µL of BER Reaction buffer, 5µL of beacon, and 5µL of lysate (at 2µg/uL nuclear lysate) was loaded. Each experimental sample was loaded in quadruplicate. Since each lysate was subjected to five different beacons; each lysate had 20 experimental wells. Three types of control samples were used: buffer only, buffer + lysate, and buffer + beacon. Each control sample was loaded in quadruplicate. The 'buffer only' control wells contain 25µL of BER buffer. The 'buffer + lysate' control wells contain 20µL buffer + 5µL lysate (at 2µg/uL nuclear lysate). All lysates had corresponding buffer + lysate wells. The buffer + beacon control wells contained 20µL buffer + 5µL beacon. All five beacons had corresponding buffer + beacon wells. Once all reagents were added to each well, the 96-well plate was sealed with Optical Adhesive Covers. The 96-well plate was then lightly vortexed and centrifuged at 500g for 10 seconds. The plate was then read on a StepOnePlus Real-Time PCR System, as we described previously. The plate was read in two stages: (1) The first stage consisted of maintaining 37°C and measuring fluorescence every 20 seconds for a total of 180 cycles. This represents the actual experimental data of the experiment; (2) The second stage ramps the temperature to 5 stages in succession: 60°C, 65°C, 70°C, 75°C, and 80°C. At each stage the fluorescence is measured every 20 seconds for a total of 15 cycles. The purpose of this stage is to determine the temperature of maximum fluorescence (Tmax ) for a given lysate + beacon combination. Dividing the experimental fluorescence by the Tmax for each beacon normalizes the data; it converts relative fluorescence to normalized fluorescence. Normalized fluorescence values allow inter-plate comparisons.

Analysis: Each experimental value had their corresponding 'buffer + lysate' and the 'buffer + beacon' values subtracted from the value. The 'buffer only' value was then added back. The baseline was then adjusted by subtracting cycle 15 from each experimental value (ignoring all values before cycle 15) yielding baseline adjusted values. Fluorescence in the second stage was then analysed to determine the $T_{max}$ temperature. The baseline adjusted values were then divided by the fluorescence at $T_{max}$. This yields normalized fluorescence values in quadruplicate. The four values were then averaged and plotted against time.

**Flow cytometry**

B16F10, B16F10 Cas9, B16F10 ΔUNG, B16F10 ΔSTING, YUMM1.7 (parental), and YUMM1.7 ΔUNG (polyclonal) cells were treated with 5 µM ATRi (AZD6738) or DMSO (0.05%), or transfected with control DNA or dU-modified DNA oligos (IDT), or treated with IFN-gamma (0.1-1.0 ng/mL) for the indicated durations. Cells were harvested via scraping, pelleted, resuspended in flow cytometry staining (FCS) buffer (Invitrogen), and aliquoted to 96-well round-bottom plates. Cells were blocked in 100 µL 5% normal mouse serum (Invitrogen) in 1x PBS for 10 min on ice, stained with MHC-I and PD-L1 or isotype control antibodies (all 1:100 in 100 µL FCS buffer) for 1 h on ice in the dark, and stained with efluor780 fixable viability dye (1:3000 in 100 µL 1x PBS) for 10 min on ice in the dark, with FSC buffer washes between staining steps. Cells were analyzed live or were fixed in 175 µL FluoroFix (BioLegend) for 2 h at room temperature in the dark and washed in FSC buffer prior to analyses. Single-stained OneComp eBeads (Invitrogen) were used as compensation controls for MHC-I and PD-L1. A single-stained mix of live and heat-killed (30 sec at 95°C) cells were used as the compensation control for efluor780. Acquisition was performed with a 4-laser CytoFLEX (Beckman Coulter) and analyses were performed in FlowJo V10.

For in vivo immune profiling experiments, B16F10 Cas9 or ΔUNG tumors were harvested from mice at the indicated time points. Corresponding splenocytes were used for single color controls, fluorescence-minus-one (FMO) controls, and for general gating. Tumors were weighed prior to processing. Single cell suspensions were generated from tumors and spleens. Tumor tissue (≤ 250 mg) was minced and then digested in Collagenase IV Cocktail (approximately 2 mL per 100 mg tumor) containing 3.2 mg/mL Collagenase IV, 1 mg/mL DNAse I, 2 mg/mL Soybean Trysin Inhibitor (all Worthington) for two 15 minute incubations at 37°C, with periodic agitation, and tituration steps between incubations and after the second incubation. Tumor homogenate was smashed through a 70 µm cell strainer (Corning) using the rubber plunger of a syringe and the filter was rinsed with 1x PBS. Tumor samples were centrifuged at 350 x g for 5 minutes and pellets were resuspending in complete D-MEM media. Spleens were mechanically dissociated between frosted glass slides and filtered through 70 µm cell strainers. Erythrocytes were lysed in 150 mM NH4Cl, 10 mM NaHCO3, 0.1 mM EDTA pH 8.0. for 10 sec (tumors) or 30 sec (spleens). Cell suspensions were counted with a Scepter 3.0 (Millipore) and seeded at 1.2-2 x $10^6$ cells (equivalent number within an experiment) in 96-well round bottom plates for blocking and staining as follows: Fc receptors were blocked for 10 min at 4°C with 0.5 µg anti-CD16/32 antibody

(TruStain FcX Plus, BioLegend) in FSC buffer (Invitrogen), cells were stained with antibodies to surface antigens (in FSC buffer) for 15 min at 4°C, dead/dying cells were stained with eFlour780 fixable viability dye (1:2000-3000, Invitrogen) or LIVE/DEAD fixable near-IR dead cell stain (1:1000, Invitrogen) in 1x PBS for 10 minutes at 4°C, samples were fixed and permeabilized in eBioscience Fixation/Permeabilization reagent (Invitrogen) for 15 min at room temperature, and when performing nuclear (Ki67, Foxp3) or intracellular cytokine (IFN-γ, TNF-α, IL-17) staining, samples were stained for 45 min at room temperature in eBioscience 1x Permeabilization Buffer (Invitrogen) containing antibodies to nuclear/intracellular proteins. Brilliant Stain Buffer Plus (BD Biosciences) was added to antibody cocktails containing multiple Brilliant Violet dye conjugates to prevent polymer dye-dye interactions. True-Stain Monocyte Blocker (BioLegend) was added to surface antibody cocktails containing PE-Cy7 conjugates to prevent non-specific binding of monocytes/macrophages to the tandem dye. For measurement of cytokine-producing $CD8^+$ and $CD4^+$ T cells, prior to staining, cells were stimulated for 4 h with 1x eBioscience Cell Stimulation Cocktail (plus protein transport inhibitors) (Invitrogen), which contains PMA and ionomycin, in complete D-MEM media. Unstimulated controls were treated for 4 h with 1x eBioscience Cell Protein Transport Inhibitors cocktail (Invitrogen) in D-MEM media. Since PMA/ionomycin stimulation induces internalization of CD3 and the CD8/CD4 co-receptors, staining for CD3, CD4, CD8, and CD45 was performed post-fixation/permeabilization during intracellular cytokine staining. Uncompensated data were collected using a BD LSRFortessa 4-laser cytometer and BD FACSDiva software. Compensation and data analyses were performed in FlowJo V10 software. Single stained spleen samples with matching unstained cells or single stained OneComp eBeads (Invitrogen) were used for single color compensation controls. Fluorescence-minus-one (FMO) controls were used, where appropriate, to empirically determine gating. Gating strategies are shown in Figures S9-S13.

**Cell-survival analysis**

Cell survival analysis was performed using cell titer-Glo® 2.0 assay (Promega, Fittsburgh, USA) according to instructions provided by the company. Briefly, tumor cells were seeded in 96-well microtiter plates and kept at 37°C in a 5% CO2 incubator. The next day, cells were treated with the indicated concentration of AZD6738, AZD0156, or DMSO. Cells were exposed to cell titer-Glo® 2.0 reagent for 10min at RT, and the obtained luminescence value was measured by a plate reader (BioTek Synergy/LX multimode reader). The relative cell survival analysis was calculated in relation to DMSO treated samples.

**RNA extraction and qPCR**

Cells were seeded overnight in a 35 mm dish. The next day they were treated with inhibitors or transfected with oligos for indicated times. At the time of harvesting, cells were washed once with PBS, and Trizol (in a 1:3 ratio) was then added to the cellular suspension. Total RNA was extracted using the direct zol RNA kit (Zymo Research Corp, Irvine) according to the manufacturer's instructions. Total RNA (typically 500ng) was reverse transcribed using the Lunascript mastermix (NEB) and diluted to add 12 ng cDNA in each reaction. Each PCR reaction was run as technical replciates using 5µl of Luna Universal qPCR mastermix (NEB), 1 µl of each 2.5 µM primer, and 3 µl of diluted cDNA (total 12 ng cDNA). qPCR was performed on a Quantstudio3 (Thermo). The PCR program was run according to the manufacturer's suggestions, and quantification was done using the Quantstudio software. Subsequent analysis of qPCR ct values was performed in Prism (Graphpad), normalizing to GAPDH expression.

**RNA-seq and data analysis**

RNA was isolated from untreated or treated cells, as described above. The isolated RNA was submitted to the Novogen Advancing Genomics facility (Sacramento, CA, USA) for RNA sequence analysis.

For RNA sequence analysis, the original data file from a high-throughput sequence platform was transferred to sequence read by CASAVA base recognition and stored in FASTQ format files. Raw data were further processed by removing adapter reads, reads with uncertain nucleotide constitute more than 10%. Read alignment was performed using HISAT2 software using the mouse reference genome. FPKM (Fragments per kilobase of transcripts sequence per millions base pairs sequenced) was used to quantify the abundance of transcripts or genes. Differential gene expression analysis was performed using DEseq2 (for biological replicates) or edgeR (for no biological replicates) software with FDR correction by the Benjamini-Hochberg procedure with threshold [log2(foldchange)]>=1 & padj <=0.05. Group pathway analysis was performed using ClusterProfiler software for Gene Ontology analysis (Gene ontology analysis (http://www.geneontology.org/ ). Gene set enrichment analysis (GSEA 4.3.2) was utilized to rank based identification of most enriched pathway between groups. GO terms with padj <0.05 are significantly enriched, and the most significant terms were selected for display.

**Immunoblotting**

Cells were lysed in 50mM Tris-HCl, pH 7.5, 150mM NaCl, 50mM NaF, 0.5% Tween-20, 1% NP40, and protease inhibitors for 20 min on ice. Lysates were cleared by centrifugation, and supernatants were collected. The soluble protein amount was estimated using protein assay kit (Biorad), and 15 ug protein was mixed with 2X Laemmi buffer (Biorad) and boiled for 10min.

Proteins were resolved in 4-12% Bis-tris, transferred to 0.4 µm nitrocellulose membrane (Bio-rad), and immunoblotted.

**Cytosolic DNA extraction and qPCR**

6 million cells were resuspended in 600 µL of cytosolic extraction buffer containing 150mM NaCl, 50mM HEPES pH=7.5 (Corning), and 25 µg/mL digitonin (Cayman Chemical). Whole-cell homogenate was divided into two aliquots of 250 µL. The first aliquot was further diluted in 250 µL of cytosolic extraction buffer, rotated end-over-end rotator for 10 minutes at 4°C, and centrifuged for 15 min at 1000xg. Four hundred microliters of supernatant was recovered and centrifuged for 10 min at 20,000xg to recover 300 µL of cytosolic fraction. DNA was isolated from the 300 µL cytosolic fraction and the second 250 µL whole-cell homogenate aliquot using Quick-DNA Miniprep Plus Kit (Zymo Research) according to the manufacturer's instructions. A sample containing 0.1% Triton-X100 in cytosolic extraction buffer (to permeabilize nuclear and mitochondrial membranes) was processed as described above and used as a positive control. Quantitative PCR was performed on 4 µL of cytosolic fractions or Triton-X100 fractions and 4 µL of whole-cell extracts diluted 1:5. DNA was amplified using the CFX Connect Real-time PCR system (Bio-Rad) and the PowerUp™ SYBR™ Green Master Mix (Applied Biosystems) following the manufacturer's instructions. The nuclear genes Gusb and Tubb5 were amplified (primers designed with the Integrated DNA Technologies (IDT) web tool) under the following conditions: 5 min at 95°C, 40 cycles of 10 sec at 95°C, and 7 sec at 62° C. Amplification was followed by a melting curve program: 15 sec at 95°C, 1 min at 70°C, ramping to 95°C while continuously monitoring fluorescence. Each sample was assayed in triplicate. The relative cytosolic fraction or Triton-X100 fraction DNA levels were normalized to whole-cell DNA levels using the delta-delta CT method.

**Animal experiments**

Experiments were performed in accordance with protocols approved by the University of Pittsburgh Animal Care and Use Committee. Female 6-8 week old C57BL/6 mice and athymic nude mice were purchased from Jackson Laboratories. B16F10 Cas9 or ΔUNG ($2.5 \times 10^5$ cells) in DMEM were subcutaneously injected into the right hind flank of 8–10-week-old mice. Tumors were measured with digital calipers on the indicated days, and volumes calculated as volume = (length x width2)/2. For studies comparing the rates of growth of B16F10 Cas9 versus B16F10 ΔUNG in nude mice or C57BL/6 mice, only tumors that estabslihed were included in measurements (i.e. tumors that did not take were excluded). For the study involving anti-PD-L1 therapy, mice were injected intraperitoneally (i.p.) with 100 µg anti-PD-L1 antibody (clone 10F.9G2, BioXCell inVivoPlus) diluted in 100 µL inVivoPure pH 6.5 Dilution Buffer (BioXCell), every 3 days for 6 doses, starting on day 2 after tumor cell injection on day 1. Control mice were injected i.p. with 100 µL inVivoPure pH 6.5 Dilution Buffer. Since treatment began the day after tumor cell injection, all tumors, including those that did not take or that completely resolved, were included in the study group measurements. The tumor endpoint was reached when tumor volume exceeded 1000 $mm^3$ or the tumor ulcerated. When multiple tumors within a group reached their endpoint, the entire study group was deemed to have reached the experimental endpoint. Mice that were euthanized due to tumor ulceration prior to the experimental endpoint were excluded from the reported tumor volumes.

**CRISPR/cas-9 mediated B16 Sting KO cells**

B16F10 STING knockout cells were generated using gene knockout kit V2 (Synthego). Multi-guide specific to mouse *TMEM173* was transfected into cells using the company's standard protocol. Briefly, the RNP complex was prepared by adding sg:RNA and cas9 in a 1.3:1 ratio, and the transfection solution was prepared using Lipofectamine CRISPRMAX. After 5-10 incubation at room temperature, these two solutions were mixed and kept for 30min at RT. Next, B16F10 ($1.5*10^3$ cells/well) were reverse transfected with the mixed solution (RNP complex and transfection solution) and transfected cells were seeded into 24 wells. Twenty-four hours later, transfection media was replaced with fresh media and kept for continue culture in the incubator. Sting knockout cells was validated by qPCR analysis and Western blot analysis.

**Quantification of nucleosides incorporated into the genome**

Cells were resuspended in 200 µL PBS. Proteinase K, RNase A, and buffer were added in quantities recommended by the PureLink™ Genomic DNA Mini Kit (Invitrogen, K182002) protocol. Samples were incubated overnight in a 55°C water bath. Genomic DNA was purified using the PureLink™ Genomic DNA Mini Kit (Invitrogen, K182002). Columns were spun at 13,000 rpm for 1 minute. DNA was eluted with RNase/DNase free water at 13,300 rpm for 1 minute. 5 µg DNA was digested with 2 µl RNase A (Thermo Scientific, EN0531), 1 µl RNase H (New England BioLabs, M0297), 3 µl Hind III (Thermo Scientific, FD0504), 3 µl EcoRI (Thermo Scientific, FD0274), 3 µl Bam HI (Fisher, FD0054) in 25 ul RNase H buffer (New England BioLabs, B0297), 25 µl Fast Digestion Buffer (10X, Thermo Scientific, B64) and 138 µl RNase/DNase free water overnight at 37 °C. Digested DNA was purified using the GeneJET PCR Purification Kit (Thermo Scientific, K0702). The column was washed with 750 µl Wash Buffer. DNA was eluted with RNase/DNase free water. Technical replicates consisting of 500 ng samples of DNA were further digested into single nucleosides using DNA Degradase Plus (Zymo Research, E2021) overnight at 37 °C. Samples were boiled on a 96 °C heat block for 10 min and then put on ice for 2 minutes. A second DNA Degradase Plus (Zymo Research, E2021) digestion was performed overnight at 37 °C. To quantitate DNA constituent base composition, a fit-for-purpose LC-MS/MS assay was implemented on a 1290 Infinity II Autosampler and Binary Pump (Agilent) and a SCIEX 6500+ triple quadrupole mass spectrometer (SCIEX). Chromatographic separation was conducted on an Inertsil ODS-3 (3 μm x 100 mm 2.1 mm) reverse phase column (GL Sciences) at ambient temperature with a gradient mobile phase of methanol and water with 0.1% formic acid. MRM transitions of all analytes and isotopic internal standards were monitored to construct calibration curves. We were able to quantitate 1 rN per 20,000 bases from 1 mg DNA.

**Micronuclei and cGAS staining**

B16F10 wild-type and ΔUNG were seeded at 2,300 cells per well on an 8-well Culture Slide (Corning #354118). The following day, cells were treated with vehicle or 5 μM AZD6738 for 48 h. After the treatment, the cells were fixed with 2 % PFA for 10 min at RT and washed with PBS. Cells were blocked with blocking buffer (2% BSA and 0.1% Triton X-100 in PBS) for 30 min at RT. This was followed by incubation with primary antibody overnight and secondary antibody for 1 h. Both primary and secondary antibodies were diluted in blocking buffer. The cells were mounted using Fluoroshield with DAPI.

**In vitro BER for dU+ micronuclei**

MEF wild-type and ΔUNG cells were seeded at 0.1 x $10^6$ on a 35 mm dish with glass bottom (MatTek, #P35G-1.5-14-C). The next day, cells were treated with vehicle or 5 μM AZD6738 for 24 h. After washing with PBS (0.2% Tween-20 in PBS), cells were incubated with freshly prepared cytoskeletal buffer (100 mM NaCl, 300 mM sucrose, 10 mM PIPES pH 6.8, 3 mM $MgCl_2$, and 0.5% Triton X-100) on ice for 5 min. This was followed by washes with PBST and fixation with 2% PFA for 10 min at RT. For permeabilization, cells were incubated with 0.25 % Triton X-100 for 10 min at RT. After washing with PBST, cells were labeled with Cy3-dUTP using a reaction that involved base-excision repair. Briefly, the reaction cocktail contained 5 U Uracil DNA-Glycosylase, 15 U Endonuclease IV, 4 U Bst full-length polymerase, 68 U Taq ligase, 500 μM NAD+, 200nM dATP, 200nM dCTP, 200nM dGTP, and 200nM Cy3-dUTP. The cells were incubated with reaction cocktail in a humid chamber for 1 h at 37°C. Cells were washed with PBST and mounted using Fluoroshield with DAPI.

**Imaging Analysis**

Cells were imaged using Nikon ECLIPSE Ti2 microscope. Micronuclei present in perinuclear region were counted using ImageJ and each field of view consisted of cells > 50. For Cy3-dUTP staining, foci that colocalized with micronuclei were regarded as positive. Nuclei that appeared apoptotic were excluded from analysis.
